# Metabolic control by the Bithorax Complex-Wnt signaling crosstalk in *Drosophila*

**DOI:** 10.1101/2024.05.31.596851

**Authors:** Rajitha-Udakara-Sampath Hemba-Waduge, Mengmeng Liu, Xiao Li, Jasmine L. Sun, Elisabeth A. Budslick, Sarah E. Bondos, Jun-Yuan Ji

**Affiliations:** Department of Biochemistry and Molecular Biology, Tulane University School of Medicine, Louisiana Cancer Research Center, 1700 Tulane Avenue, New Orleans, Louisiana 70112, USA; Lewis-Sigler Institute of Integrative Genomics, Princeton University, Princeton, New Jersey 08540, USA; Department of Medical Physiology, College of Medicine, Texas A&M University Health Science Center, College Station, Texas 77843, USA

**Keywords:** Wnt/Wingless signaling, the Bithorax Complex, lipid homeostasis, adipocyte heterogeneity, *Drosophila*

## Abstract

Adipocytes distributed throughout the body play crucial roles in lipid metabolism and energy homeostasis. Regional differences among adipocytes influence normal function and disease susceptibility, but the mechanisms driving this regional heterogeneity remain poorly understood. Here, we report a genetic crosstalk between the *Bithorax Complex* (*BX-C*) genes and Wnt/Wingless signaling that orchestrates regional differences among adipocytes in *Drosophila* larvae. Abdominal adipocytes, characterized by the exclusive expression of *abdominal A* (*abd-A*) and *Abdominal B* (*Abd-B*), exhibit distinct features compared to thoracic adipocytes, with Wnt signaling further amplifying these disparities. Depletion of *BX-C* genes in adipocytes reduces fat accumulation, delays larval-pupal transition, and eventually leads to pupal lethality. Depleting Abd-A or Abd-B reduces Wnt target gene expression, thereby attenuating Wnt signaling-induced lipid mobilization. Conversely, Wnt signaling stimulated *abd-A* transcription, suggesting a feedforward loop that amplifies the interplay between Wnt signaling and *BX-C* in adipocytes. These findings elucidate how the crosstalk between cell-autonomous *BX-C* gene expression and Wnt signaling define unique metabolic behaviors in adipocytes in different anatomical regions of fat body, delineating larval adipose tissue domains.

## Introduction

Adipocytes distributed in different parts of the body differ in their function and regulatory mechanisms. These differences include different metabolic profiles, which are essential for maintaining lipid metabolism and energy homeostasis in the whole organism. Disruptions of these processes are closely associated with diseases such as obesity, diabetes, cardiovascular diseases, and a variety of cancers ^1–6^. Notably, the specific locations of excess fat accumulation can pose distinct health vulnerabilities: visceral fat accumulated around internal organs significantly increases the risk of diseases compared to the subcutaneous fat stored in the lower body ^6–8^. Thus, it is crucial to gain a deeper understanding of the signaling pathways and developmental processes that regulate adipocyte diversity across different regions of the body.

*Drosophila* is a prominent model organism that serves as an excellent experimental system for studying the *in vivo* regulation of metabolism, especially lipid metabolism, mechanisms facilitating inter-organ communications, and disease modeling ^9,10^. The *Drosophila* fat body functions similarly to mammalian adipose tissue in regulating lipid homeostasis ^9,11,12^. In the *Drosophila* larva, fat body consists of a monolayer of adipocytes aligned along the anteroposterior axis of the larval body ^13^. This experimental model can be used to study the signaling pathways and developmental processes that regulate adipocyte diversity and lipid homeostasis. Although the fundamental metabolic processes – glycolysis, the TCA cycle, and lipid metabolism – are nearly identical in both *Drosophila* and mammals ^14,15^, major events in lipid metabolism such as adipogenesis, lipogenesis, and lipolysis occur at different developmental stages in *Drosophila*. Adipogenesis occurs exclusively during late embryogenesis, while lipogenesis, lipolysis, and fatty acid β-oxidation are predominantly active during the larval and pupal stages ^13–15^. This distinct temporal separation contrasts with the intricately interconnected and complex nature of these processes in mammals, making *Drosophila* an ideal model to isolate and analyze these distinct metabolic programs.

Wnt signaling has been shown to negatively regulate adipogenesis ^16,17^, but its role in regulating lipogenesis and lipolysis is less clear. Several reports have suggested that the expression of lipogenic enzymes such as ATP citrate lyase (ACLY), acetyl-CoA carboxylase (ACC1), fatty acid synthase (FASN), and stearoyl-CoA desaturase (SCD1) is reduced in β-catenin deficient mouse adipocytes ^18^. ChIP-seq analysis has also revealed the enrichment of Tcf7l2/TCF4 at lipogenic genes in cultured adipocytes, suggesting a direct stimulation of the expression of these lipogenic factors by Wnt signaling. Further, inhibiting GSK3 using LiCl reduced the expression of *FASN1* in the liver of juvenile turbot and decreased the triglyceride levels in plasma ^19^. Despite this, it is still unclear how Wnt signaling regulates lipid homeostasis. Our previous work in *Drosophila* larval adipocytes revealed that Wnt/Wingless (Wg) signaling directly represses the expression of Wnt target genes involved in *de novo* lipogenesis, fatty acid β-oxidation, and lipid droplet-associated proteins, thereby inhibiting lipogenesis and stimulating lipolysis and lipid mobilization ^20^. This mechanism provides a single signal that ceases storage and stimulates the mobilization of stored fat *in vivo*.

The evolutionarily conserved homeobox (Hox) genes are fundamental to establishing and conferring anteroposterior segmental differences in body plan elaboration in almost all animal lineages ^21,22^. In *Drosophila*, the Hox genes are organized into two clusters: the *Antennapedia Complex* (*ANT-C*) and the *Bithorax Complex* (*BX-C*). The *ANT-C* controls the segment identity of the anterior region of the fly’s body (head and part of the thorax), while the *BX-C* is responsible for the segment identity of the posterior region (thorax and abdomen). The *BX-C* cluster contains three Hox genes: *Ultrabithorax* (*Ubx*), *abdominal-A* (*abd-A*), and *Abdominal-B* (*Abd-B*). The roles of these BX-C proteins in regulating anteroposterior segmentation and the body plan during embryogenesis have been extensively studied ^22–24^, however their postembryonic roles in different tissues are less understood. Although the BX-C proteins are present in the abdominal fat body ^25^, the functional significance of their presence is unexplored.

Here, we report the interplay between Wnt signaling and *BX-C* proteins Abd-A and Abd-B in regulating lipid homeostasis in different regions of the larval fat body. We observed that Abd-A and Abd-B, but not Ubx, are specifically expressed in the abdominal adipocytes, where they modulate Wnt target gene expression and lipid metabolism. Depletion of BX-C genes in larval adipocytes leads to reduced fat storage, delayed larval-pupal transition, and eventual pupal lethality. Additionally, Wnt signaling stimulates the transcription of *abd-A* and, to a lesser extent, *Abd-B*, forming a feed-forward loop between Wnt signaling and Abd-A and Abd-B. This regulatory network amplifies regional adipocyte differences along the anteroposterior body axis, contributing to the precise maintenance of lipid homeostasis in *Drosophila* larvae.

## Results

### Regional difference in larval adipocytes correlates to differential expression of the BX-C proteins

This study started with an intriguing observation that connected fat storage and Wnt signaling. At the third instar wandering stage, *Axin^127^* mutant larvae become transparent (Fig. 1B, compared to the control shown in Fig. 1A), and eventually died as pupae ^26^. These phenotypes are common in conditions that disrupt lipid storage, which accounts for the opacity of larvae. It had been previously established that defective Axin (Axn) activates the canonical Wnt signaling pathway, disrupting lipid homeostasis by decreasing triglyceride levels and increasing free fatty acids ^20,26^. Active Wnt signaling stimulates lipolysis and fatty acid β-oxidation while inhibiting lipogenesis via transcriptional repression of key factors involved in these processes ^20^. However, we noted that the lipid accumulation defects in *Axn^127^* mutant larvae were consistently limited to the abdominal region of the fat body, while the fat body in the thoracic region remained unaffected (Fig. 1B). This regional difference among adipocytes has not yet been described. We observe similar regional differences in the fat body using the GAL4-UAS system to activate Wnt signaling in the larval fat body, showing that the phenomenon is not an artifact of the *Axn^127^* mutant allele or some unrelated lesion on the *Axn^127^* chromosome. We further showed that depleting *Axn* using multiple fat body-specific Gal4 drivers, such as *SREBP-Gal4* (Fig. 1C compared to Fig. 1D and Fig. S1A compared to S1B), *dCg-Gal4* (Fig. S1D cf. S1E), and *r4-Gal4* (Fig. S1F cf. S1G), consistently resulted in defects in the abdominal region of the fat body, but not the thoracic region. Depleting *slmb* (*supernumerary limbs*), encoding a conserved E3 ubiquitin ligase responsible for the degradation of Armadillo (Arm, β-catenin homolog in flies) ^27^, also activated Wnt signaling and disrupted lipid metabolism in adipocytes ^20^. Again, only the abdominal region of *slmb*-depleted larvae became transparent, while the thoracic region was unaffected (Fig. S1C cf. Fig S1A). At the cellular level, larval adipocytes from either thoracic or abdominal regions of *w^1118^* (Fig. 1E-E’) or the *SREBP-Gal4* heterozygous control (Fig. 1G-G’) larvae exhibited adipocytes of uniform size and shape. However, upon activation of Wnt signaling in *Axn^127^* homozygotes, or in *Axn*-depleted larvae, lipid accumulation in the abdominal region was significantly reduced compared to the thoracic region of the fat body (Fig. 1F’ cf. Fig. 1F and Fig. 1H’ cf. Fig. 1H). Adipocytes in the thoracic region consistently display uniform sizes (Fig. 1F and Fig. 1H), whereas adipocytes in the abdominal region are predominantly smaller and interspersed with occasional large adipocytes (Fig. 1F’ and Fig. 1H’). These observations show that the larval adipocytes are segmentally distinctive, challenging the prevailing notion that larval adipocytes are uniform and homogenous. Further, the current view that Wnt acts uniformly on fat metabolism is likely more complex, as active Wnt signaling-induced fat body defects are limited to heretofore undescribed adipocyte subtypes.

**Fig. 1.**
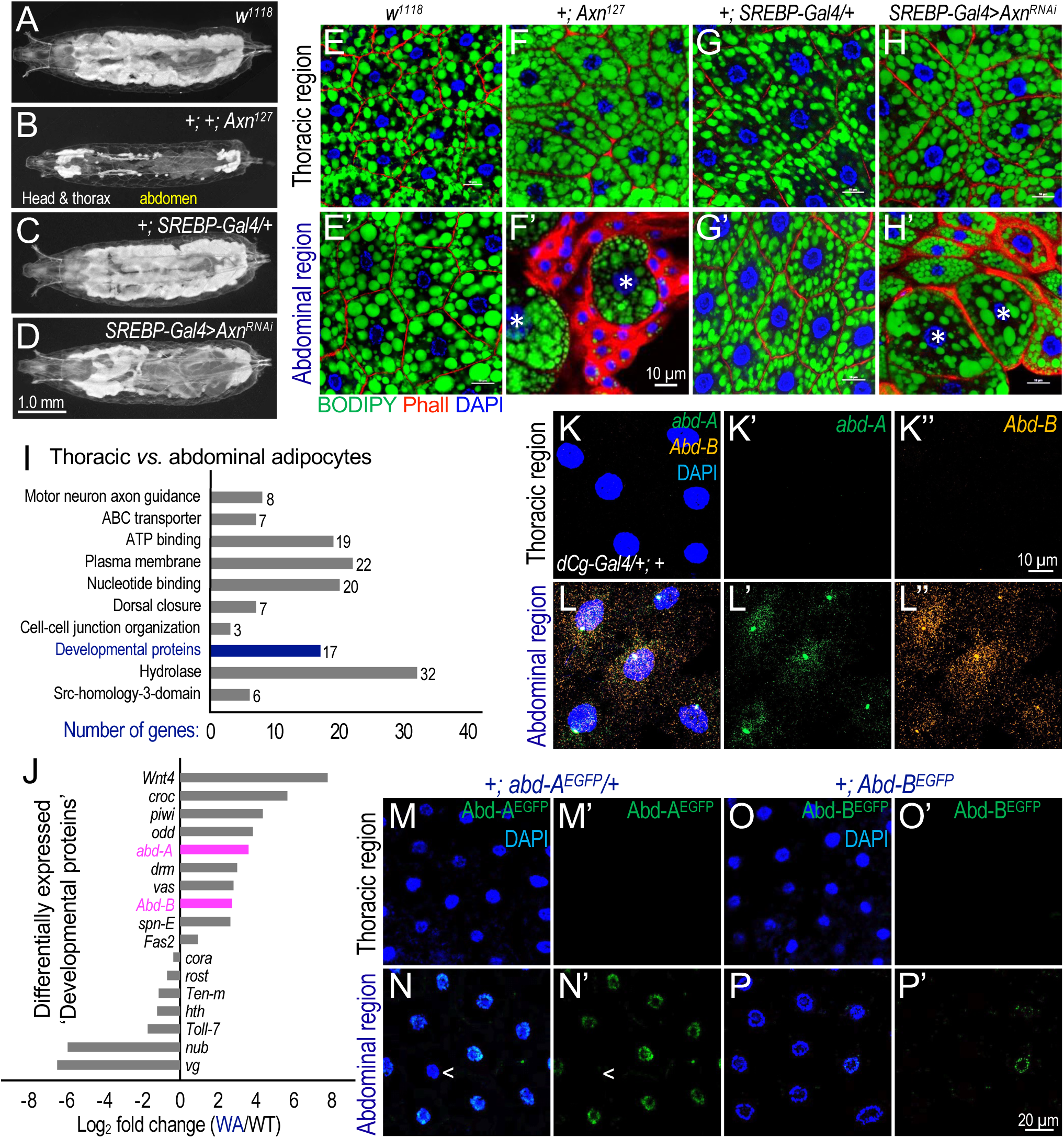
**Regional difference in *Drosophila* larval fat body correlates to differential expression of *Abd-A* and *Abd-B.*** Whole larvae images of (A) *w^1118^* (B) +; *Axn^127^* (C) *+; SREBP-Gal4/+* (D) *UAS-Axn^RNAi^/+; SREBP-Gal4/+* exhibiting the transparency in abdominal region upon activation of Wnt signaling. (E-H) Representative confocal images of larval adipocytes from larvae of indicated genotypes stained with DAPI (blue), BODIPY (green), and Phalloidin (Phall, red). Occasional large adipocytes are marked with asterisks (*). Genotypes: (E/E’) *w^1118^* (F/F’) +; *Axn^127^* (G/G’) *+; SREBP-Gal4/+* (H/H’) *UAS-Axn^RNAi^/+; SREBP-Gal4/+*. Scale bar in panel (F’) applies to images shown in E/E’-H/H’: 10μm. (I) Common pathways that are differentially regulated between thoracic *vs.* abdominal regions of fat body dissected from ‘*dCg-Gal4/+;+*’ larvae (n = 3, independent biological repeats). (J) Expression patterns of 17 Genetic factors that are categorized under ‘Developmental proteins’. *abd-A* and *Abd-B* exhibit significantly higher expression levels in the abdominal region of larval fat body compared to the thoracic region. (K/L) Hybridization chain reaction (HCR) RNA fluorescence *in situ* hybridization (RNA-FISH) imaging targeting mRNA transcripts of *abd-A* (green) and *Abd-B* (magenta) in larval adipocytes dissected from ‘*dCg-Gal4/+;+*’ control larvae. Scale bar in panle (K”) applies to images shown in K-L: 10 μm. (M-P) Expression pattern of endogenously EGFP tagged Abd-A and Abd-B in larval fat body. Genotypes: (M/M’ and N/N’) ‘*+; abd-A^EGFP^/+*’ and (O/O’ and P/P’) ‘*+; Abd-B^EGFP^*’. Occasional adipocytes that do not exhibit *abd-A* expression are marked with ‘<’. Scale bar in panel (P’) applies to images shown in M/M’-P/P’: 20 μm.

The large size of larval adipocytes and the abundant lipid droplets present technical challenges to using single-cell RNA-seq analyses to further characterize the heterogeneity of larval adipocytes. Therefore, to investigate the mechanisms underlying the differences between thoracic and abdominal adipocytes, we conducted transcriptomic analysis on groups of adipocytes following the dissection and separation of these two larval fat body regions from control ‘*dCg-Gal4/+; +*’ animals. We identified genes expressed differentially between these tissue regions and performed “Pathway” and “Gene Ontology” cluster analyses. We found significant differences in the expression levels of the genes involved in several major biological processes, including ‘Hydrolase,’ ‘Plasma membrane,’ ‘Nucleotide binding,’ and ‘Developmental proteins’ between thoracic and abdominal regions of the fat body (Fig. 1I).

Within the ‘Developmental Protein’ category, we identified 17 genes, of which 10 showed significantly higher mRNA levels in the abdominal region compared to the thoracic region (Fig. 1J). Five out of these 10 genes (*piwi*, *vas*, *spn-E*, *croc*, and *Fas2*) are expressed in gonads ^28^, which is consistent with the presence of the primordial gonads in the abdominal fat body. Removal of the primordial gonads was not possible, and we excluded these genes from further analyses. We also observed that depleting or overexpressing three other differentially expressed genes in the fat body – *Drm* (*Drumstick*) and *Odd* (*Odd-skipped*), Zinc finger transcription factors controlling blastoderm segmentation, and *Wnt4* – did not affect fat accumulation or adipocyte size (Fig. S2A-S2C). These genes were also excluded from further analyses.

Our attention was drawn to the two remaining genes, *abd-A* and *Abd-B* (Fig. 1J), because of their role in defining regions along the anteroposterior body axis. These genes encode two of the three members of the BX-C; and the third – *Ubx* – exhibit no differences in expression level. Our findings confirm previous work documenting the expression of the BX-C proteins in the abdominal fat body of *Drosophila* larvae ^25^, though the functional significance of these proteins in the fat body remains unexplored. Because the antibody used in that study could not discriminate between Abd-A and Ubx (FP3.38) ^25,29^, it was unclear whether the expression of Abd-A and Ubx was *bona fide* or an artifact. To examine Ubx expression in the fat body, we used an endogenous EGFP-tagged Ubx line generated using the CRISPR-Cas9 technique ^30^. We confirmed the expression of Ubx^EGFP^ fusion proteins in the larval central nervous cord (Fig. S4A-S4A’) and haltere discs (Fig. S4B-S4B’) ^30^. However, we did not detect any Ubx^EGFP^ protein in the entire larval fat body (Fig. S4C-S4D’). Moreover, depleting Ubx in larval fat body did not affect triglyceride levels (Fig. S5A). Therefore, our subsequent analyses focused on the function of Abd-A and Abd-B.

To analyze the differential expression of *abd-A* and *Abd-B* in individual larval adipocytes, we used multiplexed *in situ* hybridization chain reaction RNA fluorescence *in situ* hybridization (HCR RNA-FISH) imaging technology ^31^. As expected, we found mRNA levels of *abd-A* and *Abd-B* were significantly higher in the abdominal adipocytes (Fig. 1L-1L’’) compared to the thoracic adipocytes in wild-type larvae (Fig. 1K-1K’). To further analyze the expression of Abd-A and Abd-B, we genetically tagged the endogenous *abd-A* and *Abd-B* loci with EGFP using CRISPR-Cas9 (Fig. S6A and Fig. S6B). These strains were validated by sequencing. Notably, endogenous Abd-A^EGFP^ (Fig. 1M/M’ compared to Fig. 1N/N’) and Abd-B^EGFP^ (Fig. 1O/O’ compared to Fig. 1P/P’, consistent with Abd-B antibody staining Fig. S3A/A’ compared to Fig. S3B/B’) were observed in the nuclei of abdominal adipocytes but remained undetectable in the thoracic adipocytes. The expression of *abd-A* mRNA transcripts or Abd-A^EGFP^ protein within abdominal adipocytes exhibited heterogeneity, with some sporadic adipocytes exhibiting either no expression or undetectable levels of *abd-A* (Fig. 1N’). These observations reveal that the endogenous *abd-A* and *Abd-B* are expressed in abdominal adipocytes but not thoracic adipocytes.

### The differential expression of Abd-A and Abd-B defines the region-specific effects of Wnt signaling on lipid metabolism

The correlation between the region-specific effects of Wnt signaling on lipid accumulation and the differential expression patterns of Abd-A and Abd-B in abdominal adipocytes prompted us to investigate a potential interplay between BX-C proteins and Wnt signaling in shaping regional heterogeneity of adipocytes (Fig. 2A). Our hypothesis posits that altering the levels of Abd-A and Abd-B will regulate lipid metabolism, which is controlled by Wnt signaling. To test this hypothesis, we analyzed whether the ectopic expression of *abd-A* or *Abd-B* in thoracic adipocytes together with *Axn* depletion could mimic the defects observed in abdominal adipocytes by *Axn* depletion alone. In the control, *Axn* depletion (*UAS-Axn^RNAi^*/+; *SREBP-Gal4/+*) reduced fat accumulation in most adipocytes, as previously reported ^20^, accompanied by occasional large adipocytes (comparing Fig. 2C’ to Fig 2B’). In contrast, thoracic adipocytes showed no discernible effects, maintaining uniform size with a substantial accumulation of lipid droplets in each adipocyte (comparing Fig. 2C to Fig. 2B). However, the ectopic expression of wild-type *abd-A* (Fig. 2E/2E’) or *Abd-B* (Fig. 2G/G’) in this background induced heterogeneity among thoracic adipocytes (Fig. 2E and Fig. 2G). Notably, the ectopic expression of *abd-A* alone (Fig. 2D/2D’) or *Abd-B* alone (Fig. 2F/2F’) in the fat body did not alter adipocyte size compared to the control (Fig. 2B/B’). Thus, the gain of Abd-A or Abd-B in thoracic adipocytes exacerbated Wnt signaling-induced adipocyte defects, highlighting the critical role of Abd-A or Abd-B as key determinants in the effects of active Wnt signaling in larval adipocytes.

**Fig. 2.**
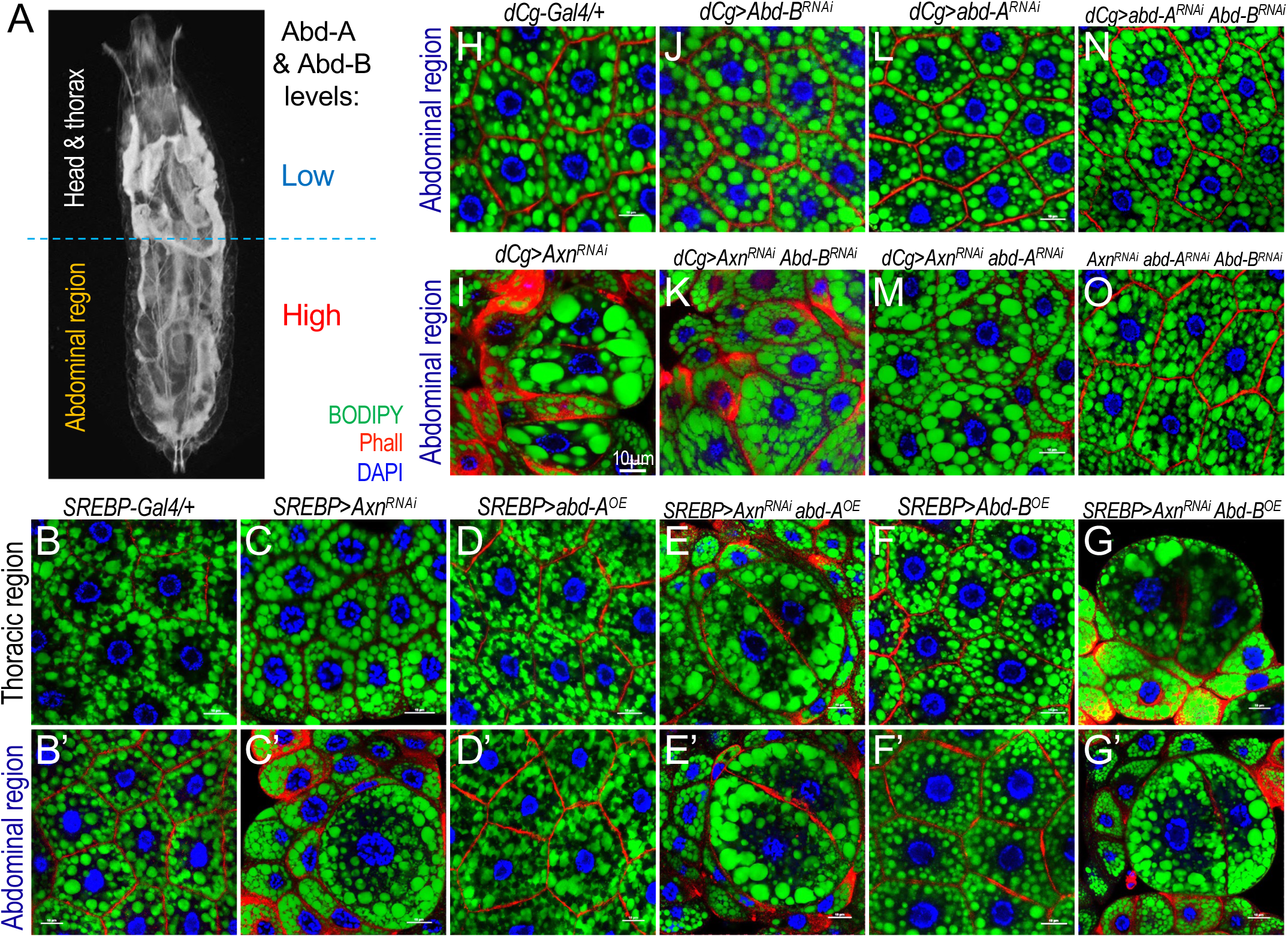
Differential expression of abd-A and Abd-B determine the region-specific activation of Wnt signaling in larval adipose tissue. (A) Schematic diagram showing transparency in the abdominal region of larval fat body upon activation of Wnt signaling and relative abundance of Abd-A and Abd-B in larval adipose tissue. (B-G) Representative confocal images of larval adipocytes from the thoracic region (B-G) and abdominal region (B’-G’) of laval fat body. Detailed genotypes: (B/B’) *+/+; SREBP-Gal4/+*; (C/C’) *UAS-Axn^RNAi^/+; SREBP-Gal4/+*; (D/D’) *UAS-Abd-A^+^/+; SREBP-Gal4/+* (OE: overexpression); (E/E’) *UAS-Abd-A^+^/UAS-Axn^RNAi^; SREBP-Gal4/+*; (F/F’) *UAS-Abd-B^+^/+; SREBP-Gal4/+*; and (G/G’) *UAS-Abd-B^+^/UAS-Axn^RNAi^; SREBP-Gal4/+*. (H-O) Confocal images of abdomical adipocytes showing significant rescue of adipocyte defects upon depleting *abd-A*, *Abd-B*, or both. Detailed genotypes: (H) *dCg-Gal4/+; +*; (I) *dCg-Gal4/UAS-Axn^RNAi^; +*; (J) *dCg-Gal4/+; UAS-Abd-B^RNAi^/+*; (K) *dCg-Gal4/UAS-Axn^RNAi^; UAS-Abd-B^RNAi^/+*; (L) *dCg-Gal4/+; UAS-Abd-A^RNAi^/+*; (M) *dCg-Gal4/UAS-Axn^RNAi^; UAS-Abd-A^RNAi^/+*; (N) *dCg-Gal4/+; UAS-Abd-A^RNAi^/UAS-Abd-B^RNAi^*; and (O) *dCg-Gal4/UAS-Axn^RNAi^; UAS-Abd-A^RNAi^/UAS-Abd-B^RNAi^*. Scale bar in panel (I) applies to all images in this figure: 10 μm.

If the effect of *Wnt* signaling on adipocyte heterogeneity in the abdominal region is dependent on Abd-A or Abd-B, then we expect that the simultaneous depletion of Axn along with Abd-A (or Abd-B) proteins would mitigate the adipocyte defects induced by active *Wnt* signaling in abdominal adipocytes. As shown in Fig. 2M, abdominal adipocytes indeed display greater uniformity in size when both *Axn* and *abd-A* were co-depleted, compared to the depletion of *Axn* alone (Fig. 2I). Knocking down *Abd-B* also reduced adipocyte heterogeneity caused by depleting *Axn* (Fig. 2K cf. Fig. 2I), albeit to a lesser extent than by *abd-A* depletion (Fig. 2M cf. Fig. 2I). The combined depletion of *abd-A* and *Abd-B* with *Axn* resulted in a robust suppression of adipocyte heterogeneity caused by *Axn* depletion (Fig. 2O cf. Fig. 2I), suggesting partial redundancy between *abd-A* and *Abd-B*. In contrast, depleting *abd-A* or *Abd-B* individually (Fig. 2L and Fig. 2J) or together (Fig. 2N) in the controls showed no discernible effects on adipocyte heterogeneity. Moreover, the transparency observed in larvae due to *Axn* depletion (Fig. S1E and S6A) was partially alleviated by co-depleting *abd-A* (Fig. S7A). A substantial rescue of the transparent larvae phenotype becomes evident when both *abd-A* and *Abd-B* were simultaneously depleted (Fig. 2O and Fig. S7A). These observations are strongly supportive of the significant roles of both *abd-A* and *Abd-B* in ameliorating fat body defects induced by active Wnt signaling.

To further explore the interplay between *abd-A*/*Abd-B* and Wnt signaling, we used the pNP vector, an efficient and multi-target transgenic RNAi system ^32^, to establish new transgenic lines enabling depletion of *abd-A* and *Abd-B* individually, as well as both simultaneously. Validation by sequencing is shown in Fig. S8A. When *abd-A* depletion occurred alongside active Wnt signaling (Fig. S8F), a pronounced reversal of fat body defects was observed compared to *Axn* depletion alone (Fig. S8E). The rescue effect became even more robust when both *abd-A* and *Abd-B* were co-depleted under conditions of active Wnt signaling (Fig. S8G cf. S8E). Controls involving the depletion of *abd-A* (Fig. S8C) or simultaneous depletion with *Abd-B* (Fig. S8D; heterozygous *r4-Gal4* control in Fig. S8B) showed no discernible effects on adipocyte heterogeneity. These observations suggest a pivotal role of Abd-A and Abd-B in modulating the effect of Wnt signaling on lipid metabolism, thereby establishing a functional link between BX-C proteins and Wnt signaling in regulating lipid homeostasis in larval adipocytes.

### The effects of depleting Abd-A and Abd-B in larval adipocytes on lipid metabolism-related gene expression

Our recent study revealed the role of Wnt signaling in regulating lipid homeostasis in larval adipocytes ^20^. However, the specific role of *BX-C* proteins in adipocyte lipid homeostasis remains unexplored to date. We observed a significant reduction in triglyceride accumulation in wandering stage larvae upon depletion of *abd-A* or the simultaneous depletion of both *abd-A* and *Abd-B* (Fig. 3A). In contrast, depletion of *Abd-B* alone resulted in marginal effects, indicating that Abd-A plays the key role in regulating lipid metabolism among the BX-C proteins. Moreover, the simultenous depletion of all three *BX-C* genes in the fat body (genotype: “*dCg-Gal4/pNP-abdA-AbdB-Ubx^RNAi^; +*”; referred to as “TKD”) resulted in a similar reduction in triglyceride accumulation to the concurrent depletion of *abd-A* and *Abd-B* (Fig. S5A, S5B). This is consistent with the very low or undetectable Ubx^EGFP^ in larval adipocytes (Fig. S4C/D). These animals exhibited a one-day delay in the larval-pupal transition (Fig. S5C). Additionally, ∼70% of TKD animals die at the pupal stage (Fig. S5E), with the remaining 30% dying within a week after eclosion. Together, these observations suggest that Abd-A, and to a lesser extent Abd-B, plays important roles in regulating lipid homeostasis in larval fat body.

**Fig. 3.**
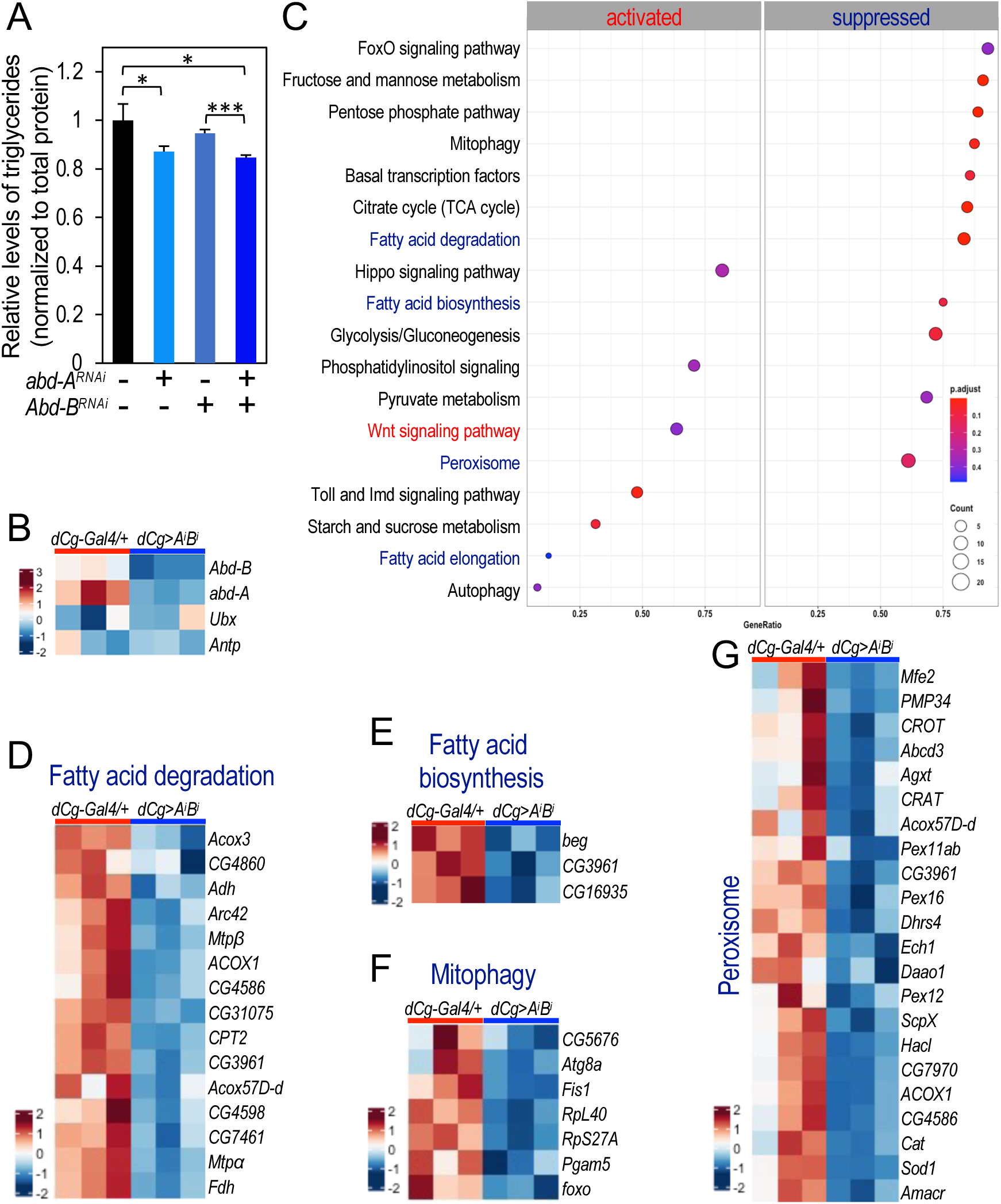
The effects of depleting Abd-A and Abd-B in larval adipocytes on lipid metabolism-related gene expression. (A) Quantification of triglyceride levels in larvae with *dCg-Gal4* driven depletion of *abd-A* and *Abd-B*. Detailed genotypes: *dCg-Gal4/+;+* (the first column); *dCg-Gal4/pNP-[abd-A]^RNAi^; +* (the second column); *dCg-Gal4/pNP-[Abd-B]^RNAi^; +* (the third column); and *dCg-Gal4/pNP-[abd-A Abd-B]^RNAi^; +* (the fourth column). (B) A heatmap of the mRNA levels of *Abd-B*, *abd-A, Ubx,* and *Antp* in ‘*dCg-Gal4/+;+*’ (*dCg/+*) and ‘*dCg-Gal4/pNP-[abd-A Abd-B]^RNAi^/ +*’ (abbreviated as ‘*dCg/A^i^B^i^*’). (C) Common pathways related to developmental signaling, lipid and carbohydrate metabolism that are significantly altered in larval fat body with the depletion of *abd-A* and *Abd-B.* (D-G) Heatmaps representing the gene expression levels of several metabolic pathways, including ‘Fatty acid degradation’ pathway (D), ‘Fatty acid biosynthesis’ pathway (E), ‘Mitophagy’ (F), and ‘Peroxisome’ pathway (G).

To systematically identify the target genes of Abd-A and Abd-B in larval adipocytes, we performed RNA-Seq analyses using dissected fat bodies from larvae depleted of both proteins, given their functional redundancy (abbreviated as ‘*AiBi*’; genotype: “*dCg-Gal4/pNP-[abdA-AbdB]^RNAi^;+*”). These analyses revealed that 1734 genes were significantly upregulated upon simultaneous depletion of *abd-A* and *Abd-B*, whereas 1249 genes were significantly downregulated. Both *abd-A* and *Abd-B* were downregulated as expected, whereas other Hox genes such as *Ubx* and *Antp* were not affected (Fig. 3B). Downregulated genes in *AiBi* showed a significant reduction in metabolic pathways related to lipid metabolism and carbohydrate metabolism (Fig. 3C). Pathways including ‘Fatty acid degradation’ (Fig. 3D), ‘Fatty acid biosynthesis’ (Fig. 3E), ‘Glycolysis/gluconeogenesis’, ‘Citrate cycle (TCA cycle)’, and ‘Pentose phosphate pathway’ were among those significantly dysregulated (Fig. 3C). Additionally, there were significant effects on genes regulating mitophagy (Fig. 3F) and peroxisomal metabolism (Fig. 3G). Furthermore, the simultaneous depletion of *abd-A* and *Abd-B* led to a prominent upregulation of signaling pathways such as Hippo, Wnt, MAPK, and TGF-β. These data indicate that Abd-A and Abd-B play important regulatory roles in regulating cellular lipid and carbohydrate metabolism, while also significantly influencing developmental signaling pathways.

### Abd-A and Abd-B modulate the expression of Wnt target genes in larval adipocytes

Our genetic analyses revealed that co-depleting *abd-A* and *Abd-B* reversed fat accumulation defects induced by active Wnt signaling in larval adipocytes (Fig. 2H-2O). To dissect the underlying mechanism, we conducted RNA-seq analyses and compared the abdominal fat body from *Axn*-depleted larvae and the control (*dCg-Gal4/+;+*), as Wnt-derived fat body defects were limited to the abdominal region. The ‘Wnt signaling pathway’ exhibited the expected upregulation, while many genes related to metabolic pathways, particularly ‘Fatty acid metabolism,’ and carbohydrate metabolism-related pathways such as ‘Glycolysis/Gluconeogenesis’, ‘Citrate cycle (TCA cycle)’, and ‘Pentose phosphate pathway’ showed significant downregulation (Fig. 4A). Notably, direct target genes of Wnt signaling, such as *Notum*, *Fz3*, and *nkd*, were significantly upregulated by activated Wnt signaling (Fig. 4B).

**Fig. 4.**
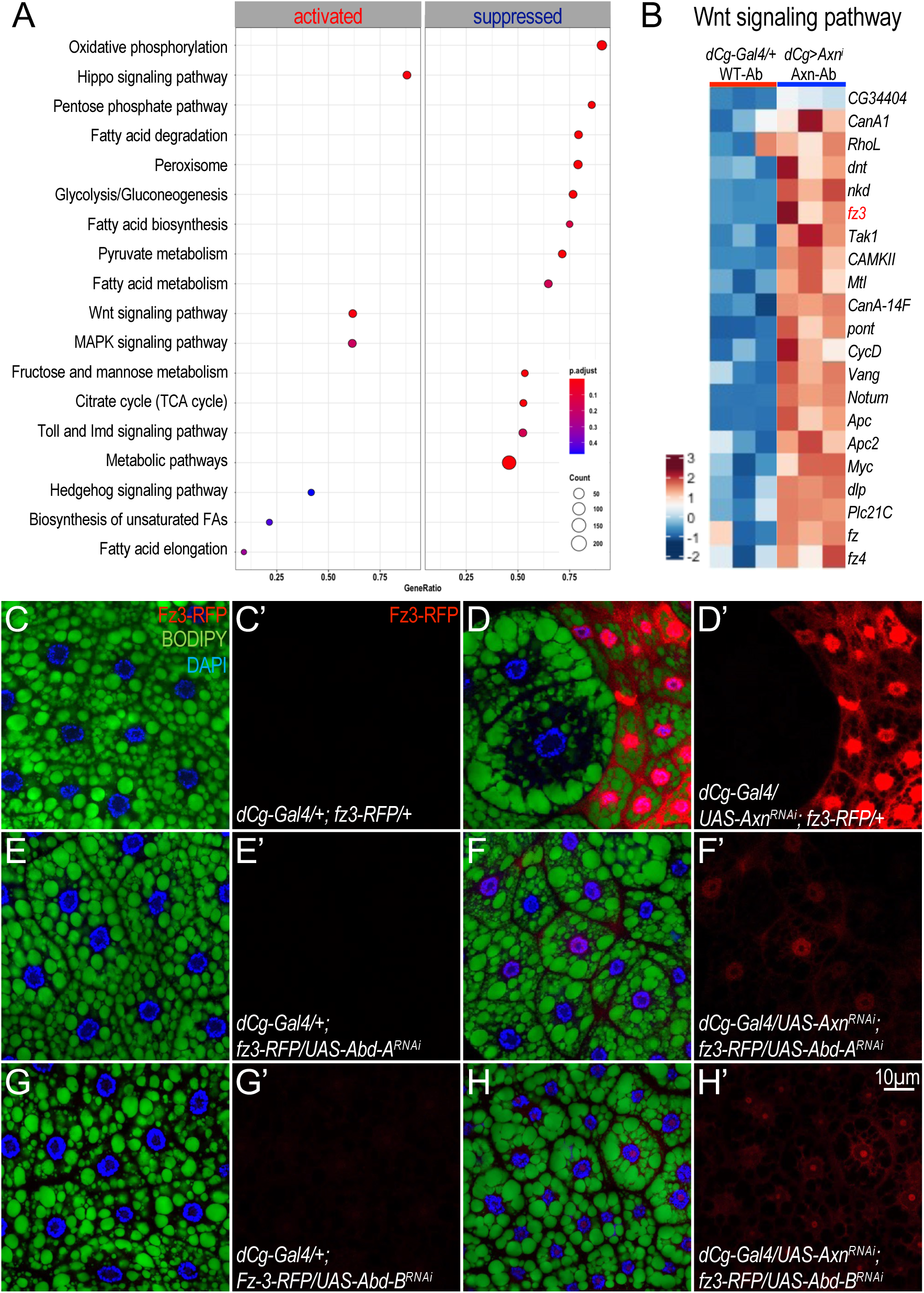
Abd-A and Abd-B are required for the expression of Wnt-activated target genes in larval adipocytes. (A) Common pathways related to developmental signaling, lipid and carbohydrate metabolism that are significantly altered in the abdominal region of larval fat body with the depletion of *Axn* (*dCg-Gal4/Axn^RNAi^; +*) in comparison to the control (*dCg-Gal4/+;+*) (B) A heatmap representing the gene expression levels of the ‘Wnt signaling pathway’ in samples from ‘WT-Ab’ (‘Wildtype Abdominal’ region of ‘*dCg-Gal4/+;+*’ larvae) *vs*. ‘Axn-Ab’ (‘Axn Abdominal’ region of ‘*dCg-Gal4/Axn^RNAi^;+*’ larvae). *fz3* is highlighted in red as it was used in the subsequent analysis. (C/C’-H/H’) Representative confocal images of abdomial asipocytes stained with BODIPY (green) and DAPI (blue). *dCg-Gal4* driven depletion of *abd-A* and/or *Abd-B* together with *Axn*, under the *fz3-RFP* background. Detailed genotypes: (C/C’) *dCg-Gal4/+; fz3-RFP/+*; (D/D’) *dCg-Gal4/UAS-Axn^RNAi^; fz3-RFP/+*; (E/E’) *dCg-Gal4/+; fz3-RFP/UAS-Abd-A^RNAi^*; (F/F’) *dCg-Gal4/UAS-Axn^RNAi^; fz3-RFP/UAS-Abd-A^RNAi^*; (G/G’) *dCg-Gal4/+; Fz-3-RFP/UAS-Abd-B^RNAi^*; and (H/H’) *dCg-Gal4/UAS-Axn^RNAi^; fz3-RFP/UAS-Abd-B^RNAi^*. Scale bar in panel (H’): 10 µm.

Given that active Wnt signaling can either positively or negatively regulate the transcription of different target genes ^20^, we sought to analyze the effect of Abd-A and Abd-B on Wnt signaling-regulated target gene expression at the cellular level. Using the *fz3-RFP* reporter as a readout for Wnt-activated genes ^33^, we observed no detectable expression in the control (Fig. 4C/4C’). However, *Axn* depletion in abdominal adipocytes resulted in high *fz3-RFP* expression in small adipocytes with active Wnt signaling, contrasting with undetectable levels in adjacent large adipocytes with low Wnt activity (Fig. 4C/4C’ cf. Fig. 4D/4D’). This is identical to our previous observations of the behavior of *nkd* ^20^, another known direct target of the Wnt signaling pathway ^20^.

To test the role of Abd-A and Abd-B in Wnt-activated target gene expression, we co-depleted them with *Axn*. Co-depletion effectively abolished the effects of *Axn* depletion on *fz3-RFP* expression. Specifically, when *Axn* was co-depleted with *abd-A* (Fig. 4F/4F’ cf. Fig. 4D/4D’) or *Abd-B* (Fig. 4H/4H’ cf. Fig. 4D/4D’), there was a substantial reduction in *fz3-RFP* expression, rescuing adipocyte heterogeneity within the abdominal fat body. However, depleting either *abd-A* or *Abd-B* alone did not cause any significant effect on *fz3-RFP* expression (Fig. 4E/4E’ and Fig. 4G/4G’). These results suggest that Abd-A and Abd-B may play a permissive role in the expression of Wnt signaling-activated genes in abdominal adipocytes.

We next asked whether Abd-A and Abd-B also regulated the expression of Wnt-repressed genes. As mentioned earlier, our pathway analyses revealed that many lipid and carbohydrate metabolism-related pathways were downregulated upon activation of Wnt signaling (Fig. 4A). Specifically, the expression levels of key regulators in fatty acid biosynthesis, including *FASN1* (encoding Fatty acid synthase 1), *bgm* (*bubblegum*, encoding a long-chain-fatty-acid-CoA ligase), (Fig. 5A) and fatty acid degradation, including *CPT2* (encoding carnitine palmitoyltransferase 2) and *Acox3* (Acyl-CoA oxidase 3), were significantly reduced (Fig. 5B). Additionally, we observed that lipid droplet associated proteins (LDAPs) such as *Lsd1* (encoding Lipid storage droplet-1) and *Lsd2* (encoding Lipid storage droplet-2) are significantly downregulated by active Wnt signaling (Fig. 5C), consistent with our previous findings ^20^.

**Fig. 5.**
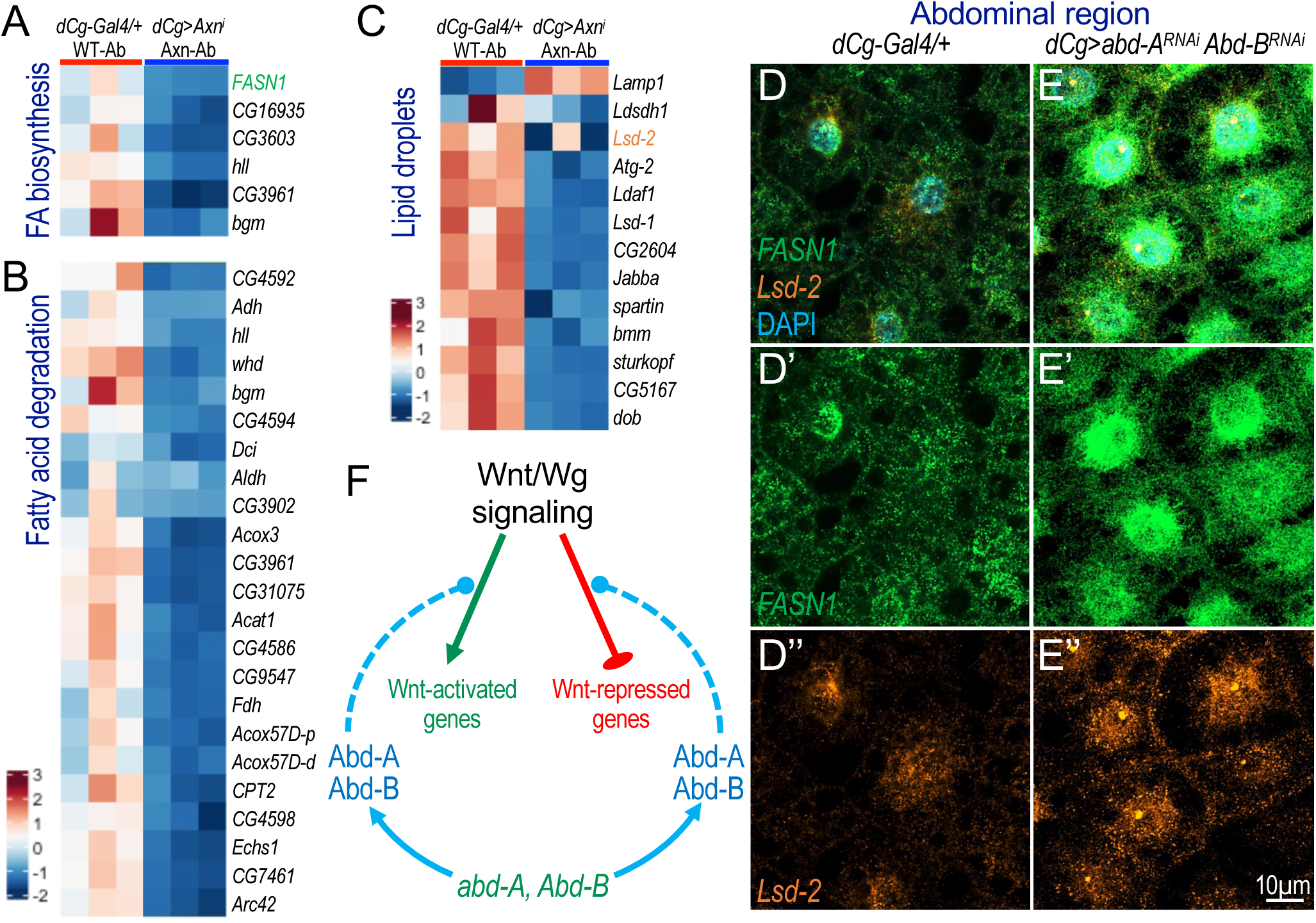
Abd-A and Abd-B are required for the expression of Wnt-repressed target genes in larval adipocytes. (A-C) Heatmaps representing the gene expression levels of the ‘Fatty acid biosynthesis’ (A), ‘Fatty acid degradation’ (B), and ‘Lipid particle’ (C) pathways in WT-Ab (‘Wildtype Abdominal’ region of ‘*dCg-Gal4/+;+*’ larvae) *vs*. Axn-Ab (‘Axn Abdominal’ region of ‘*dCg-Gal4/Axn^RNAi^;+*’ larvae). *FASN1* in panel A and *Lsd-2* in panel C are highlighted in green and orange, respectively, as those genes were used in the subsequent analyses. (D-D”’/E-E”’) Representative confocal images showing the mRNA transcripts of *FASN1* (green) and *Lsd-2* (orange) in adipocytes detected by the HCR RNA-FISH assay. Genotypes: (D-D’’) *dCg-Gal4/+* and (E-E’’) *dCg>Abd-[A+B]^RNAi^*. Scale bar in panel E’’ applies to all the images: 10 µm. (F) The model figure illustrates that Abd-A and Abd-B are required for the expression of Wnt-target genes such as *fz-3, FASN1*, and *Lsd-2*.

To further investigate the effect of depleting *abd-A* and *Abd-B* on Wnt-repressed genes, we performed HCR RNA-FISH analysis using specific probes for *Lsd2* and *FASN1*. As shown in Fig. 5E/5E’/E’’, depleting *abd-A* and *Abd-B* led to a significant increase in expression levels of both *Lsd2* and *FASN1*, compared to the control (Fig. 5D-D’’). These observations suggest that Abd-A and Abd-B are required for the expression of Wnt-repressed genes (Fig. 5F). Together, these findings reveal the significant roles of *abd-A* and *Abd-B* in modulating the expression of both Wnt-activated and Wnt-repressed target genes. The generality of this conclusion is further tested below.

### Wnt signaling potentiates the expression of *abd-A* and *Abd-B* in larval adipocytes

Similar to other crucial genes that control complex developmental processes, the *abd-A* and *Abd-B* loci are characterized by large intronic regions that may integrate complex developmental cues and signaling inputs. Upon analyzing our CUT&RUN (Cleavage Under Targets & Release Using Nuclease) assay using *dTCF^EGFP^* wing imaginal discs ^20^, we observed multiple *dTCF^EGFP^* binding peaks enriched within gene bodies of both the *abd-A* (Fig. 6A) and *Abd-B* loci (Fig. S9A and S9B). To test if Wnt signaling regulates the endogenous transcription of *abd-A* and *Abd-B*, we analyzed mRNA transcripts of *abd-A* and *Abd-B* using multiplexed HCR RNA-FISH. In fat bodies from the control larvae, the expression levels of *abd-A* and *Abd-B* were higher in the abdominal adipocytes (Fig. 6C-C’’), and were low or undetectable in the thoracic adipocytes (Fig. 6B-B’’). As we have reported recently, the smaller adipocytes display a higher Wnt activity upon activating Wnt signaling ^20^. As shown in Fig. 6E, we observed that Wnt signaling activation significantly increased *abd-A* expression in the abdominal adipocytes, especially in cells with higher Wnt activity (Fig. 6E’), in contrast to the thoracic adipocytes (Fig. 6D’). These results suggest that *abd-A* is directly regulated by Wnt signaling, leading to enhanced *abd-A* transcription in response to active Wnt signaling.

**Fig. 6.**
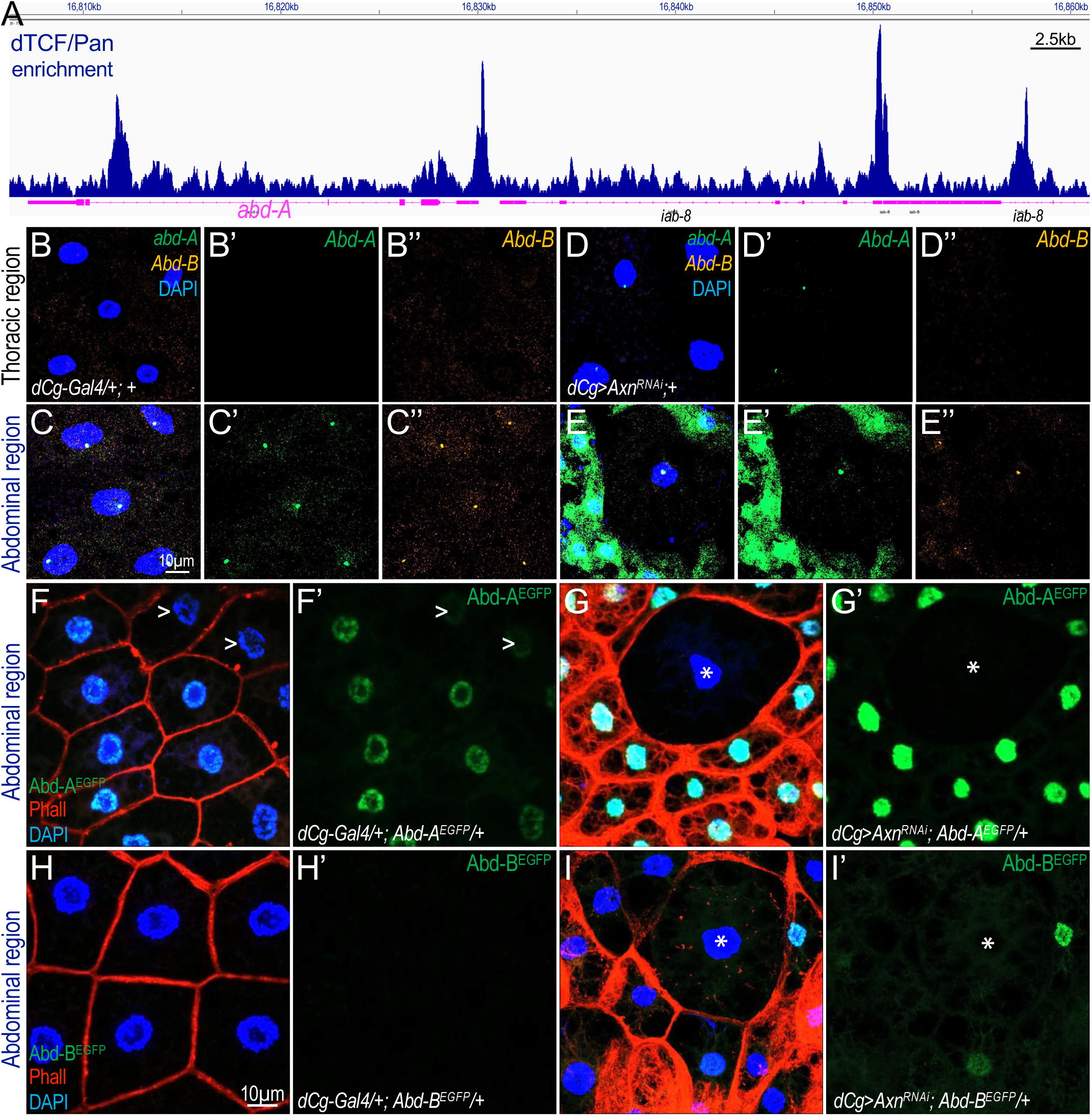
Wnt signaling directly regulates the expression of *abd-A* and *Abd-B* in larval adipocytes. (A) The screenshot displays the genomic track visualized via the Integrative Genomics Viewer (IGV) browser, illustrating dTCF/Pan binding peaks detected at the *abd-A* and a portion of the *iab-8* loci through CUT&RUN anssay performed on wing discs. The y-axis is autoscaled, with different isoforms of these genes collapsed and shown in magenta. Scale bar: 2.5kb. (B/B”-E/E”) Representative confocal images showing the mRNA transcripts of *abd-A* (green) and *Abd-B* (magenta) in thoracic adipocytes (B-B’’, D-D’’) and abdominal adipocytes (C-C’’, E-E’’), as detected by the HCR RNA-FISH assay. Scale bar in panel (C) applies to all the images shown in B-E: 10 µm. Genotypes: (B-B”/C-C’’) *dCg-Gal4/+; +*; and (D-D”/E-E’’) *dCg-Gal4/UAS-Axn^RNAi^; +.* (F/G) Depleting *Axn* under Abd-A^EGFP^ background. Genotypes: (F/F’) *dCg-Gal4/+; Abd-A^EGFP^/+* (control); and (G/G’) *dCg-Gal4/UAS-Axn^RNAi^; Abd-A^EGFP^/+.* Adipocytes with lower levels of Abd-A expression are marked with ‘>’, and adipocytes with low Wnt activity are marked with ‘*’ (asterisk). (H-I’) Depleting *Axn* under Abd-B^EGFP^ background. Genotypes: (H/H’) *dCg-Gal4/+; Abd-B^EGFP^/+* (control); and (I/I’) *dCg-Gal4/UAS-Axn^RNAi^; Abd-B^EGFP^/+.* Adipocytes with low Wnt activity are marked with ‘*’ (asterisk). Scale bar in panel H’ applies to F-I: 10 µm.

To evaluate the effect of Wnt signaling on Abd-A and Abd-B protein levels, we activated Wnt signaling in backgrounds where Abd-A or Abd-B were tagged with EGFP at their endogenous loci. In the control, Abd-A^EGFP^ was significantly higher in the abdominal adipocytes (Fig. 6F/F’) compared to the thoracic region (Fig. S10A/A’), with noticeable heterogeneity. Activating Wnt signaling consistently increased Abd-A^EGFP^ levels in abdominal adipocytes, particularly in small Wnt-active adipocytes (Fig. 6G/G’), while no obvious effects were observed in thoracic adipocytes (Fig. S10B/B’). Large adipocytes, however, exhibited notably lower expression levels of Abd-A^EGFP^ (Fig. 6G’). These observations provide further evidence for a permissive role of Abd-A in facilitating Wnt activation in larval adipocytes.

In contrast, endogenous Abd-B^EGFP^ expression was low and undetectable in most larval adipocytes of the *dCg-Gal4* heterozygous control (Fig. 6H/H’; the thoracic region is shown in Fig S10C/C’). However, active Wnt signaling also increased Abd-B^EGFP^ expression in small abdominal adipocytes (Fig. 6I/I’; the thoracic region is shown in Fig. S10D/D’). Consistent effects were observed using ectopically tagged Abd-A^EGFP^ and Abd-B^EGFP^ lines generated with re-engineered BAC clones to introduce fluorescent tags attached to specific protein isoforms ^34^ (Fig. S10E/E’ cf. S10F/F’ and Fig. S10G/G’ cf. S10H/H’). These observations suggest that active Wnt signaling stimulates the transcription of *abd-A*, with a similar but weaker effect on *Abd-B*, in larval adipocytes.

### Wnt targets, including Abd-A, are predominantly expressed in the abdominal region of the larval fat body

The results presented above suggest that both Abd-A and Abd-B positively regulate Wnt signaling (Fig. 2), which in turn directly stimulates the expression of *abd-A* and *Abd-B* (Fig. 6), these findings collectively reveal a simple model of the crosstalk between the BX-C and Wnt signaling (Fig. 7A). Given that the expression of *abd-A* and *Abd-B* in the abdominal adipocytes is significantly higher than in thoracic adipocyte (Fig. 1J), we decided to further analyze the interplay between BX-C and Wnt signaling in defining the regional difference among the larval adipocytes. Specifically, we performed RNA-seq experiments using dissected thoracic and abdominal region fat bodies from *Axn* depleted: samples from thoracic fat body of the control larvae are designated as ‘WT-Th’ for ‘wildtype thoracic’, samples from abdominal fat body are designed as ‘WT-Ab’ for ‘wildtype abdominal’, while samples from *Axn^RNAi^* thoracic fat body are called ‘Axn-Th’ for ‘*Axn^RNAi^* thoracic’, and samples from *Axn^RNAi^* abdominal fat body is designated ‘Axn-Ab’ for ‘*Axn^RNAi^* abdominal’ (Fig. 7B). Our RNA-seq analysis revealed that *Axn* is indeed depleted in the entire *Axn^RNAi^* fat body, including both ‘Axn-Th’ and ‘Axn-Ab’ samples compared to the control (Fig. 7C). As expected, the mRNA levels of both *abd-A* and *Abd-B* are higher in the abdominal adipocytes (‘WT-Ab’) than then thoracic adipocytes (‘WT-Th’), while Wnt signaling activation induced by depleting Axn significantly increased the levels of *abd-A* in abdominal adipocytes (‘Axn-Ab’) than then thoracic adipocytes (‘Axn-Th’) (Fig. 7C), while the effects on *Abd-B* expression level was marginal (Fig. 7C).

**Fig. 7.**
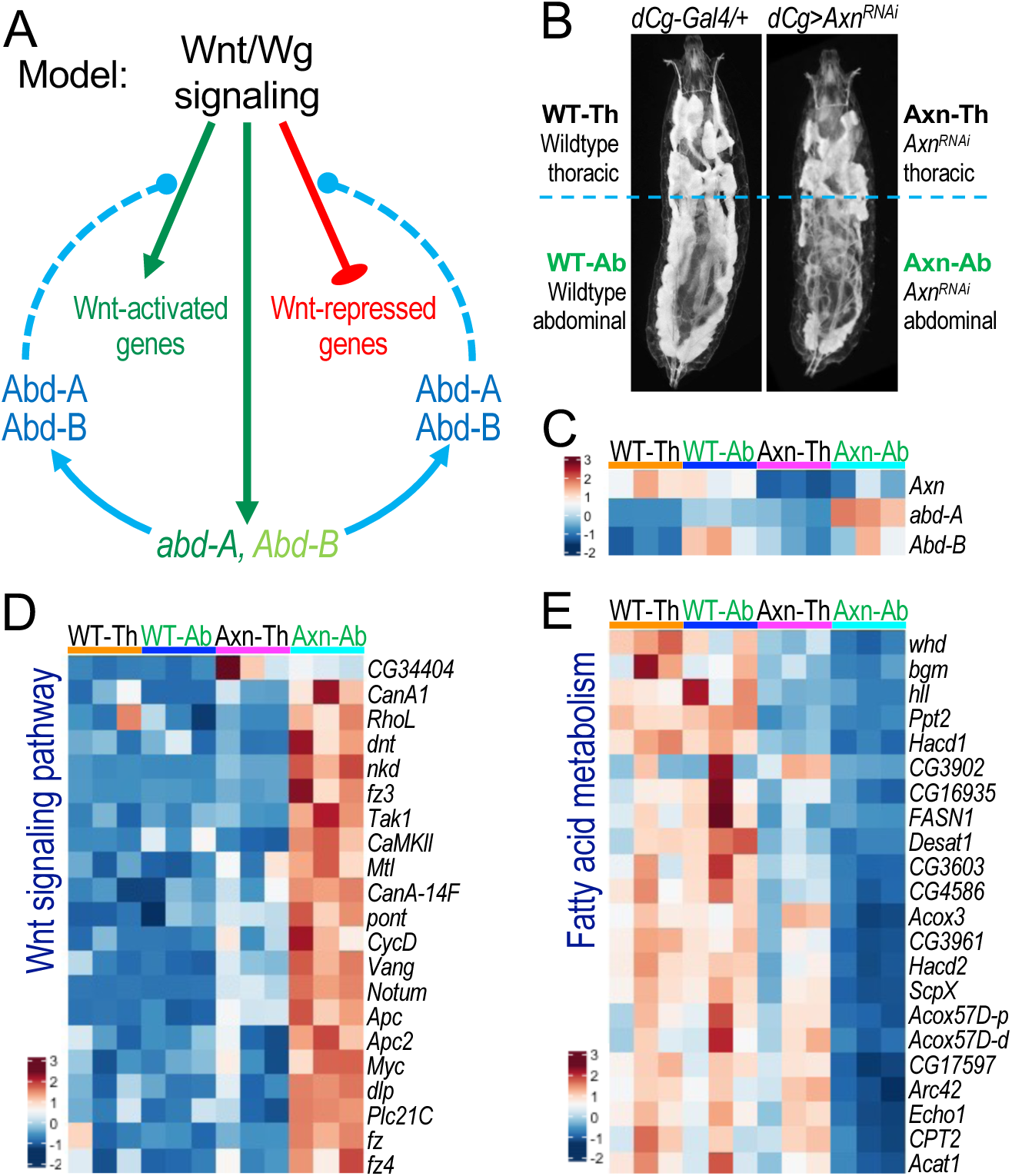
Crosstalk between Wnt signaling and Abd-A/Abd-B defines the regional difference of the larval fat body. (A) The proposed model illustrates the crosstalk between Wnt signaling and Abd-A/Abd-B: Wnt signaling may directly regulate the expression of *abd-A* and, to a lesser extent, *Abd-B*, while Abd-A and Abd-B proteins are required for both the activation and repression of Wnt target gene expression. (B) Whole larvae images showing the distinct regions used for RNA-seq analysis. Thoracic and abdominal regions of ‘*dCg-Gal4/+;+*’ (control) larva are labeled as ‘WT-Th’ and ‘WT-Ab’, respectively. Thoracic and abdominal regions of ‘*dCg-Gal4/UAS-Axn^RNAi^; +*’ larva are labeled as ‘Axn-Th’ and ‘Axn-Ab’, respectively. (C) A heatmap of the mRNA levels of *Axn*, *Abd-A* and *Abd-B* in ‘WT-Th’, ‘WT-Ab’, ‘Axn-Th’, ‘Axn-Ab’ samples. (D) A heatmap representing the gene expression levels of ‘Wnt signaling pathway’ showing that Wnt-activated gene expression is prominent in ‘Axn-Ab’, while ‘Axn-Th’ resembles ‘WT-Th’ and ‘WT-Ab’. (E) A heatmap representing the gene expression levels of ‘Fatty acid metabolism’ showing that Wnt-repressed gene expression is significantly reduced in ‘Axn-Ab’, while ‘AT’ resembles ‘WT-Th’ and ‘WT-Ab’.

Our analysis of a few candidate genes, such as *fz3* (Fig. 4), *FASN1*, and *Lsd-2* (Fig. 5), indicates an essential role of Abd-A and Abd-B modulating Wnt target genes that are either activated or repressed by active Wnt signaling. However, the generality of this notion remained unknown. Using the DAVID GO analysis of the RNA-seq data, we identified a set of Wnt-activated genes termed ‘Wnt signaling pathway’, including known Wnt-targets such as *fz3*, *nkd*, *Notum*, *CycD*, and *Myc* etc (Fig. 7D). As expected, the expression of these Wnt target genes in control fat body, both ‘WT-Th’ and ‘WT-Ab’ was low; in contrast, when Wnt signaling was activated by depleting *Axn*, Wnt-activated genes, including *fz3*, were significantly higher in abdominal adipocytes (‘Axn-Ab’ samples), while their expression in the thoracic adipocytes (‘Axn-Th” samples) remained low and resembles both the thoracic and abdominal adipocytes from the control (‘WT-Th’ and ‘WT-Ab’ samples) (Fig. 7D). These results support the permissive role of Abd-A in regulating the expression of a battery of Wnt activated genes in larval adipocytes (Fig. 7A).

Our recent work has identified a set of Wnt signaling-repressed genes encoding factors that play critical roles in regulating both lipogenesis and lipid mobilization ^20^, which are categorized under ‘fatty acid metabolism’ pathway. In control samples, their expression showed no obvious difference between the thoracic (‘WT-Th’ samples) and abdominal (‘WT-Ab’ samples) adipocytes (Fig. 7E). However, their expression was significantly reduced in Wnt-active abdominal adipocytes (‘Axn-Ab’ samples) compared to the Wnt-active thoracic adipocytes (‘Axn-Th’ samples). These data suggest that Abd-A is also required for active Wnt signaling to repress the expression of its target genes involved in lipid metabolism, including *FASN1*, *whd* (*CPT1*), *CPT2*, and others (Fig. 7E), in larval adipocytes.

These findings collectively suggest that Abd-A and Abd-B play a permissive role in regulating the expression of both Wnt-activated and Wnt-repressed genes in larval adipocytes. This effect is predominantly evident in abdominal adipocytes, where the endogenous *abd-A* and *Abd-B* are expressed (Fig. 7A). The presence of endogenous Abd-A and Abd-B in abdominal adipocytes makes them responsive to Wnt signaling-induced lipid mobilization. Conversely, adipocytes lacking *abd-A* and *Abd-B* expression, whether in the abdominal or thoracic regions, fail to activate or repress Wnt target genes. Consequently, this leads to large adipocytes in thoracic adipocytes and occasional abdominal adipocytes despite *Axn* depletion (Fig. 7C).

## Discussion

This study reveals the intricate interplay between Wnt signaling and *BX-C* proteins Abd-A and Abd-B, shaping adipocyte heterogeneity and lipid homeostasis in *Drosophila* larvae. Abd-A and Abd-B play a permissive role in modulating Wnt target gene transcription, while conversely, Wnt signaling also potentiates *abd-A* and *Abd-B* expression in abdominal adipocytes (Fig. 7A). This genetic interact architecture provides a mechanism for the origin of adipocyte heterogeneity, important for regulating lipid homeostasis in *Drosophila* larval adipocytes.

### The role of Abd-A and Abd-B in regulating lipid metabolism

Previous research has documented the expression of the *BX-C* proteins in the abdominal fat body of *Drosophila* larvae using immunostaining techniques ^25^, yet the functional significance of these proteins in adipocytes remains unknown. Our triglyceride measurements of larvae with *abd-A* and *Abd-B* depleted, specifically in adipocytes (‘*AiBi*’), revealed a substantial reduction in lipid accumulation. Consistent with this observation, our transcriptomic analyses of larval adipocytes depleted *abd-A* and *Abd-B* (‘*AiBi*’) revealed significant downregulation of metabolic pathways, including carbohydrate metabolism and fatty acid metabolism. Further analyses revealed significant impacts of depleting *abd-A* and *Abd-B* on metabolic processes such as the ‘pentose phosphate pathway’, ‘TCA cycle’, ‘fatty acid degradation’, ‘fatty acid degradation’ and ‘fatty acid biosynthesis’, among others. These findings show that Abd-A and Abd-B regulate lipid and carbohydrate metabolism *in vivo*, underscoring their physiological significance in larval adipocytes.

It is currently unclear whether the transcription of these lipid metabolism-related genes is directly or indirectly regulated by Abd-A and Abd-B in larval adipocytes. Addressing this question requires conducting ChIP-seq analyses, albeit with several technical challenges to overcome. Firstly, the use of ChIP-grade antibodies specific to Abd-A and Abd-B is imperative. The monoclonal FP3.38 antibody commercially available from the Developmental Studies Hybridoma Bank (DSHB) cannot discern Ubx from Abd-A ^25,29^, and whether the monoclonal antibody for Abd-B (1A2E9) available from the DSHB is suitable for ChIP assay is uncertain. EGFP-tagged Abd-A and Abd-B strains present an intriguing avenue for ChIP-seq or similar CUT&RUN analyses. Secondly, the presence of abundant lipid droplets and a limited number of larval adipocytes poses major technical challenges for both ChIP-seq and CUT&RUN assays. Our previous attempts using CUT&RUN assay to analyze dTCF/Pan binding sites with purified nuclei from dissected larval fat bodies yielded uninformative results due to suboptimal signal-to-noise ratios ^20^. Lastly, considering the critical role of Abd-A and Abd-B in various biological contexts, conducting the CUT&RUN assay in other tissues, such as the central nervous system ^35^, or developmental stages, such as the late embryonic stage, to infer their binding sites in larval adipocytes may prove problematic. Addressing these technical obstacles will require additional efforts and future technical breakthroughs.

The Hox family of transcription factors can function as either transcriptional activators or repressors ^36,37^. For example, Abd-A acts as a repressor, while Ubx and Abd-B function as activators, as evidenced by analysis using the *dpp674-lacZ* reporter, which is driven by a visceral mesoderm-specific enhancer of *decapentaplegic* (*dpp*), during embryogenesis ^38^. However, the ectopic expression of *abd-A* in male histoblast nest cells (HNCs) triggers *wg* expression in the seventh segment of early pupal abdominal epithelia ^39^. In the cardiac tube, Abd-A was shown to activate the transcription of *Ih* (encoding a voltage-gated ion channel) and *ndae1* (encoding a Na^+^-driven anion exchanger) during embryogenesis, while repressing *Ih* expression in the pupa ^40,41^. These findings suggest that Abd-A is capable of functioning as either a transcriptional activator or repressor. It is intriguing to explore how Abd-A modulates the expression of lipid metabolism-related genes in larval adipocytes.

### The interplay between *BX-C* proteins and Wnt signaling

Our genetic and cell biological analyses suggest that the impaired fat accumulation induced by active Wnt signaling is suppressed by depleting *abd-A* and *Abd-B*. Three potential scenarios could explain this phenomenon. Firstly, Abd-A and Abd-B might regulate lipid metabolism through their direct interactions with dTCF/Pan ^42,43^, the pivotal transcription factor downstream of the Wnt signaling pathway ^44,45^. Secondly, the effect of Abd-A and Abd-B on lipid metabolism in larval adipocytes may be direct, potentially involving shared target genes of *BX-C* proteins and Wnt signaling. As discussed earlier, it remains technically challenging to identify the direct target genes of Abd-A, Abd-B, and dTCF/Pan in larval adipocytes. Thirdly, crosstalk with additional signaling pathways, such as ‘Hippo signaling pathway’ and ‘TGF-β signaling pathway’, could also play a role, as evidenced by consistent upregulation of these pathways in the ‘*AiBi’* background ^46–48^. These scenarios are not mutually exclusive, and the complexity of these interactions warrants further investigation in future studies.

An unexpected observation is that the depletion of Abd-A and Abd-B mitigates the transcription of both Wnt-signaling activated genes and Wnt-signaling repressed genes, suggesting a permissive role of Abd-A and Abd-B proteins in regulating Wnt target gene expression in larval adipocytes. While the underlying mechanism is unknown, we speculate that certain transcription cofactors required for Wnt target gene expression, either activation or repression, may be regulated by Abd-A and Abd-B. Interestingly, it has been reported that depleting HOXB5 represses the expression of *β-catenin* and its downstream target genes, thereby significantly inhibiting tumor invasion in human non-small cell lung cancer (NSCLC) ^49^. Additionally, endogenous HOXB7 directly interacts with β-Catenin, and depletion of HOXB7 inhibits activation of the Wnt/β-catenin signaling pathway. Furthermore, HOXB8 has been shown to prevent the activation of the Wnt/β-catenin signaling pathway, thus efficiently suppressing tumorigenesis and metastasis in colorectal cancer cells ^49^. *HOXB7* is a human ortholog of the *Drosophila Ubx* gene, while *HOXB8* is the human ortholog of *abd-A* ^50^. To our knowledge, the potential role of HOXB8, HOXB7, and HOXB5 in the transcription of Wnt signaling repressed genes is yet to be determined. However, these findings indicate that the role of Abd-A and Abd-B in modulating Wnt target gene transcription could be conserved from flies to humans.

Another intriguing observation is the potentiation of *abd-A* and *Abd-B* transcription by active Wnt signaling (Fig. 6). The presence of multiple dTCF/Pan binding sites revealed by the CUT&RUN assay indicates that Wnt signaling may directly stimulate the transcription of *abd-A* and *Abd-B*. However, this effect is exclusively detectable in adipocytes where the transcription of *abd-A* and *Abd-B* is already underway, *i.e*., in abdominal adipocytes. This suggests that active Wnt signaling does not instruct or initiate the transcriptional activation of *abd-A* and *Abd-B* genes but rather synergizes with other regulatory factors to further amplify their transcription. Interestingly, the *HoxB9* gene, an orthologous gene of *Abd-B* ^50^, was reported to be a direct target of WNT/TCF signaling in mice ^51^, indicating potential conservation in Wnt signaling in regulating HOX gene expression.

These mechanisms, coupled with exclusive expression of Abd-A and Abd-B in the abdominal adipocytes, offer a compelling explanation for why the activation of canonical Wnt signaling by depleting either *Axn* or *slmb* in larval adipocytes results in distinct effects specifically in abdominal adipocytes, with minimal effect on thoracic adipocytes (Fig. 7A). Interestingly, occasional large adipocytes are consistently observed in abdominal adipocytes with active Wnt signaling ^26^. These large adipocytes lack the expression of Abd-A and Abd-B and behave similarly to thoracic adipocytes in response to Wnt signaling. Thus, our model (Fig. 7A) also offers a mechanistic explanation for this adipocyte heterogeneity within abdominal adipocytes. Furthermore, our transcriptomic and cell biological analyses reveal the key roles of *abd-A* and *Abd-B* in regulating lipid homeostasis and the transcription of Wnt target genes, with their expression being potentiated by active Wnt signaling (see below).

### Heterogeneous expression of Abd-A and Abd-B defines adipocyte heterogeneity

Due to its crucial role in establishing the anterior-posterior segmental body plan during embryogenesis, the transcriptional regulation of the *BX-C* locus has been extensively studied in the past decades ^21,52,53^. Once established, the expression status of *BX-C* genes is maintained during postembryonic stages by the combined regulation of *PcG* and *TrxG* proteins ^54–57^. However, experimental evidence for the involvement of *PcG* and *TrxG* proteins is currently lacking.

In *Drosophila* larvae, a cohesive sheet of 2100-2500 adipocytes is embedded along the anterior-posterior body axis of larvae ^13^. Previous studies by Rizki and colleagues have suggested differences among larval adipocytes based on variations in autofluorescent metabolites, such as kynurenine and other tryptophan metabolites, as well as pteridines ^13^. Also, other studies suggest that specific regions provide positional cues for the functional specialization of fat cells, facilitating the development of biochemically diverse areas within the larval/adult fat body ^58^. Notably, the endogenous *abd-A* and *Abd-B* are heterogeneously expressed in larval fat body (this work), further extending the previous report that *BX-C* proteins are differentially expressed in larval fat body ^25^. Contrary to the widely held view of larval fat bodies as homogeneous, our RNA-seq analyses of thoracic and abdominal fat body, alongside studies by Rizki, Pimpinelli, and their colleagues ^13,25^, reveal substantial heterogeneity. This study also reveals that adipocyte heterogeneity is significantly manifested by the activation of Wnt signaling. Beyond their difference in regulating lipid homeostasis, the additional functional relevance of adipocyte heterogeneity in different body regions warrants further investigation, employing improved analytical tools and approaches.

Our HCR analyses show that active Wnt signaling predominantly stimulates higher Abd-A expression in Wnt-active small adipocytes, while the large adipocytes show moderate effects in the expression of Abd-A (Fig. 6E’). Similar observations were made for EGFP-tagged Abd-B expression (Fig. 6I’ and Fig. S10I’), albeit less convincing due to the low or undetectable levels of Abd-B protein in larval adipocytes. Integrating these observations with our genetics data, which imply a permissive role of Abd-A and Abd-B in regulating Wnt target gene expression, suggests that low levels of Abd-A and Abd-B in large adipocytes mitigate Wnt activity. Therefore, the heterogeneous expression of *abd-A* and *Abd-B* in the fat body is fundamental to adipocyte heterogeneity, and dissecting the genetic circuits that control their heterogeneous expression in larval adipocytes will advance our understanding of *BX-C* gene regulation beyond embryogenesis.

### Adipocyte heterogeneity in evolution

Adipocytes play critical roles in regulating lipid homeostasis and energy metabolism in metazoans, and dysregulated adipocyte function is associated with diseases such as obesity, diabetes, and various cancers. There are three types of adipocytes in mammals: white, brown, and beige adipocytes ^59–61^. White adipocytes primarily store triglycerides, brown adipocytes specialize in thermogenesis, while beige adipocytes possess characteristics of both. Despite extensive studies in mouse models, the evolutionary and developmental origins of these different types of adipocytes remain poorly understood. Investigating their evolutionary origin may offer insights into their adaptive significance to the overall fitness of metazoans, while elucidating the developmental origin of different adipocytes may reveal key regulatory factors and signaling pathways involved in adipogenesis during embryogenesis.

In addition to the different types of adipocytes, their distribution within the body also varies with sex and development, correlating with disease susceptibilities ^6^. Accumulating evidence has linked visceral obesity to increased risks of metabolic diseases and systematic metabolic dysfunction ^6^. Notably, lipolysis in visceral adipocytes is less sensitive to insulin suppression than in leg adipocytes in humans ^62^. In our study, abdominal adipocytes are particularly sensitive to Wnt signaling-inducted lipolysis due to the specific expression of Abd-A (and Abd-B) in these adipocytes. Depleting Abd-A, Abd-B, or Axn altered the expression of genes involved in several signaling pathways, including Hedgehog, Hippo, MAPK, Toll and Imd signaling pathways (Fig. 3C and Fig. 4A). Perhaps the ligands for these signaling pathways are secreted from other tissues or organs into body cavities, functioning as signaling molecules for communications between adipose tissue and other organs. It will be interesting to determine whether adipocytes in different parts of the fat body display distinct sensitivity to these signaling pathways similar to Wnt signaling.

Exploring adipocyte diversity in invertebrates may advance our understanding of evolutionary adaption. While genes related to lipogenesis and lipolysis are present even in unicellular organisms like *Saccharomyces cerevisiae* and *Candida parapsilosis* ^63–65^, insects represent the first occurrence of dedicated adipocytes for fat storage ^66^. This distinguishes them from lower multicellular animals like nematodes, where lipid droplets are stored in gut epithelial cells. In addition to fat storage, insect adipose tissue also serves as endocrine and immune functions in insects, similar to higher organisms ^67^. We speculate that the heterogeneous adipocytes observed in *Drosophila* may represent a primitive form of white and brown adipocytes in mammals, although the adipocyte heterogeneity in *Drosophila* larvae lacks the sophisticated functions and intricate regulations of different types of adipocytes and fat depots found in mammals ^66^.

This study explored the functional relevance of *BX-C* proteins in larval adipocytes and the underlying mechanisms defining adipocyte heterogeneity in *Drosophila* larvae. Given the conservation of key components of the Wnt signaling pathway and *BX-C* proteins in metazoans ^22–24,50,68,69^, it would be intriguing to investigate whether the interplay between Wnt signaling and BX-C proteins also regulates adipocyte diversity in mammals.

## Materials and Methods

### *Drosophila* stocks and maintenance

The flies used in this study were maintained at 25°C and fed with a standard diet comprising cornmeal, molasses, and yeast medium. The *Axn^127^/TM6B* and *UAS-Axn^RNAi^* lines were described previously ^20,26^, and genotypes of other strains are provided in Table S1. Using pNP vector-based system ^32^, we generated *UAS-abd.A^RNAi^ Abd.B^RNAi^* line (specific primers are included in Table S2), and validated this line by sequencing (Fig. S8A).

### Cell biological analyses

The methods for staining the larval fat body with BODIPY, Nile Red, DAPI, and phalloidin have been previously described ^26,70^. Following the same procedures, triglyceride level quantifications were performed using a colorimetric kit as previously described ^26,70^.

### The HCR RNA-FISH assay

The multiplexed *in situ* HCR RNA-FISH assay was conducted as described previously ^70^. The following specific probe sets were obtained from Molecular Instruments: B1-Alexa Fluor 488 amplifiers with probe sets for *abd-A* (lot number PRO337); B2-Alexa Fluor 594 amplifiers with the probe sets for *Abd-B* (lot number PRQ416). Confocal images were captured using a Zeiss LSM900 confocal microscope system, and representative images for each experiment were presented.

### RNA-seq sample preparation, library preparation, sequencing, differential gene expression analysis, and gene ontology enrichment analysis

These experiments were done using the same methods described previously ^20,70^.

### Immunocytochemistry

These experiments were done following the same protocol described previously with slight modifications ^71^. Fat bodies from third-instar larvae at the late wandering stage were used. Anti-ABD-B (1A2E9) antibody from Developmental Studies Hybridoma Bank (DSHB) was used at a concentration of 5 µg/ml. Goat anti-mouse (115-545-003) secondary antibody was used (purchased from Jackson Immunological Laboratories).

### Tagging the endogenous *abd-A* and *Abd-B* loci with EGFP using the CRISPR-Cas9 technique

To generate the endogenously *abd-A* (*CG10325*) tagged line, cassette EGFP-PBacDsRed was inserted, and ATG of *abd-A-RB* was replaced by using a guide RNA (TGACGAATTCGGGGGGGTGG[TGG]). The EGFP-PBacDsRed cassette contains EGFP and a selection marker PBacDsRed. The two homology arms (upstream and downstream homology arms; Fig. S5A) of *abd-A* were amplified by Phusion High-Fidelity DNA Polymerase (Thermo Scientific) from genomic DNA at optimized conditions, which reflects sequences of the injection strain *in vivo*. Cassette EGFP-PBacDsRed and two homology arms were cloned by ‘Golden Gate’ method into vector pUC57-Kan (Donor plasmid), followed by standard transformation protocol, colony PCR selection, and sequencing. The selection marker PBacDsRed contains 3’ PBac terminal repeats, the artificial 3xP3 promoter (three tandem copies of the Pax-6 homodimer binding site and TATA-homology of hsp70), DsRed2, SV40 3’UTR, and 5’ PBac terminal repeats. It facilitates genetic screening and can be excised by Piggy Bac transposase. Only one TTAA motif will be left after transposition. Sequence GTTAAA will encode Val Lys as a linker between EGFP and gene. ‘Excision PCR’ technique was used to excise the selection marker where forward primer (‘*Exicision-abd.A5.1*’) was designed at EGFP and reverse primer (‘*Exicision-abd.A3.1*’) was designed downstream of the homology arm. The selection marker was excised when examining the endogenous protein expression using EGFP-tagged Abd-A. PAM mutations (TGACGAATTCGGGGGGGTGG[T**<u>C</u>**G]) were made to keep the donor inactive to guide RNA incorporated into the edited genome.

The same approach was used with *Abd-B-RB* (*CG11648*) to generate endogenously EGFP-tagged Abd-B (Fig. S5B). Cassette EGFP-PBacDsRed was inserted, and ATG of *abd-B-RB* was replaced using a guide RNA (AGATGGTGCTGCTGCATGAC[GGG]). PAM site mutation was not required for this construct design. Primer sequences are included in Table S2.

### Statistical analyses

All P-values in this study were determined using one-tailed unpaired *t*-tests, and the error bars shown in the figures represent the standard deviation. Each experiment in this work was analyzed with a minimum of three independent biological replicates. * for p<0.05, ** for p<0.01, and *** for p<0.001.

## Acknowledgement

We thank Yashi Ahmed, Ingrid Lohmann, and Jian-Quan Ni for generously sharing fly strains and reagents, and Keith Maggerts and Michael Levine for their insightful comments on the manuscript. We thank the Bloomington *Drosophila* Stock Center (NIH Grant P40OD018537) for fly stocks, and the Developmental Studies Hybridoma Bank at the University of Iowa for monoclonal antibodies. This research was supported by a grant from the National Institute of Health (GM129266 to J.-Y.J.).

---

**Supplementary Fig. S1. Whole larvae images showing the depleting *Axn* or *slmb* using several fat body specific Gal4 drivers.** Note the specific reduction in abdominal fat accumulation. Detailed genotypes: (A) *+; SREBP-Gal4/+*; (B) *UAS-Axn^RNAi^ /+; SREBP-Gal4/+*; (C) *+; SREBP-Gal4/UAS-Slmb^RNAi^*; (D) *dCg-Gal4/+; +*; (E) *dCg-Gal4/UAS-Axn^RNAi^; +*; (F) *+; r4-Gal4/+*; and (G) *UAS-Axn^RNAi^ /+; r4-Gal4/+*. Scale bar in (G): 0.5 mm.

**Supplementary Fig. S2. The ectopic expression or depletion of *Wnt4* does not cause any obvious phenotypic effects in adipocytes**. (A-C) Representative confocal images of larval adipocytes from larvae of indicated genotypes stained with DAPI (blue), BODIPY (green), and Phalloidin (Phall; red). Detailed genotypes: (A) *dCg-Gal4/+; +,* (B) *dCg-Gal4/UAS-Wnt4^+^; +* (OE: overexpression); and (C)*dCg-G4/+; UAS-Wnt4^RNAi^/+.* Scale bar in (A): 10 μm.

**Supplementary Fig. S3. Antibody staining of *w^1118^* larval fat body using monoclonal antibodies raised against Abd-B**. The thoracic region (A/A’) and abdominal regions (B/B’) of wandering larvae fat body are shown separately. The red color represents Abd-B and DAPI (blue) stains nuclei. Scale bar in (B): 20 μm.

**Supplementary Fig. S4. Endogenous expression pattern of EGFP tagged Ubx (Ubx^EGFP^)**. Endogenous expression pattern of Ubx^EGFP^ in the VNC (ventral nerve cord) region (A-A’), haltere discs (B-B’), thoracic region of the larval fat body (C-C’), and abdominal region of fat body (D-D’). The green color represents Ubx and DAPI (blue) stains nuclei. Scale bar in (D’): 10 μm.

**Supplementary Fig. S5. Effects of *BX-C* gene depletion in larval fat body.** Triple knockdown of the three *BX-C* genes (“*TKD*”; genotype: ‘*dCg-Gal4/UAS-abd.A-Abd.B-Ubx^RNAi^; +*’) in the larval fat body results in a reduction of triglyceride accumulation (n.d: no significant difference). Triglyceride levels were measured using a Triglyceride Assay kit (A) and thin layer chromatography (TLC, B). Lane 1 in (B) shows the lipid standard, which is a mixture of monoglyceride (MG), diglyceride (DG), TG, and stearic acid (representing Free Fatty Acids/FFA). (C) The larval-pupal transition in *TKD* larvae is delayed. (E) The TKD aminals are pupal lethality, with the control (ctrl: ‘*dCg-Gal4/+; +*’) shown in (D). Both pupae were at day 9 AEL (after egg laying).

**Supplementary Fig. S6. CRISPR-Cas9 based EGFP-tagging of *abd-A* and *Abd-B*.** CRISPR/Cas9-mediated genome editing by homology-dependent repair (HDR) using one guide RNA and a dsDNA plasmid donor. The N termini of Abd-A (RB form) (A) and Abd-B (RB form) (B) were tagged using “EGFP-PBacDsRed” cassette, which contains EGFP and 3xP3-DsRed flanked by PiggyBac terminal repeats. Two homology arms were cloned into pUC57-Kan as donor templates for repair. Scale bar: 10 kb.

**Supplementary Fig. S7.** Whole larvae images for *dCg-Gal4* driven *abd-A* and *Abd-B* depleted larvae with or without active Wnt signaling induced by depleting *Axn.* Note that Wnt signaling-induced fat body defects in the abdominal region of the fat body are significantly rescued by co-depleting *abd-A* and *Abd-B*.

**Supplementary Fig. S8. Validation of *abd-A^RNAi^ Abd-B^RNAi^ Drosophila* strains**. (A) Sequence validation of pNP-based *abd-A* and *Abd-B* double depletion lines using Sanger sequencing. (B-G) Depleting *abd-A* and/or *Abd-B* under active Wnt signaling background using pNP vector-based system. Detailed genotypes: (B) *+; r4-Gal4/+*; (C) *_pNP_UAS-Abd-A^RNAi^/+; r4-Gal4/+*; (D) *_pNP_UAS-Abd-A^RNAi^ Abd-B^RNAi^/+; r4-Gal4/+*; (E) *UAS-Axn^RNAi^/+; r4-Gal4/+*; (F) *_pNP_UAS-Abd-A^RNAi^/UAS-Axn^RNAi^; r4-Gal4/+*; (G) *_pNP_UAS-Abd-A^RNAi^ Abd-B^RNAi^/UAS-Axn^RNAi^; r4-Gal4/+.* Scale bar in (E): 10 μm.

**Supplementary Fig. S9.** The genomics tracks display dTCF/Pan binding peaks identified in parts of *iab-8* (A) and *Abd-B* (B) loci through CUT&RUN analysis conducted on wing discs. These gene tracks are visualized using the IGV browser. Together with the genomic track shown in Fig. 6A, these genomic tracks encompass the entire *abd.A-Abd.B* genomic region. The y-axis is autoscaled, and different isoforms of those genes are collapsed and shown in magenta. Scale bars: 2.5 kb.

**Supplementary Fig. S10. Effects of Wnt signaling on *abd-A* and *Abd-B* expression in larval adipocytes.** (A-B) Thoracic adipocytes stained with DAPI (blue) and Pholl (red) in *Abd-A^EGFP^* background. Genotypes: (A/A’) *dCg-Gal4/+; Abd-A^EGFP^/+*; and (B/B’) *dCg-Gal4/UAS-Axn^RNAi^; Abd-A^EGFP^/+*. (C-D) Thoracic adipocytes stained with DAPI (blue) and Pholl (red) in *Abd-B^EGFP^* background. Genotypes: (C/C’) *dCg-Gal4/+; Abd-B^EGFP^/+*; and (D/D’) *dCg-Gal4/UAS-Axn^RNAi^; Abd-B^EGFP^*. (E-F) Abdominal adipocytes stained with DAPI (blue) and Pholl (red) in *Abd-A^EGFP^*background. Genotypes: (E/E’) +; *r4-Gal4/Abd-A^EGFP^*; and (F/F’) *UAS-Axn^RNAi^/+; r4-Gal4/Abd-A^EGFP^*. (G-H) Abdominal adipocytes stained with DAPI (blue) and Pholl (red) in *Abd-B^EGFP^*background. Genotypes: (E/E’) +; *r4-Gal4/Abd-B^EGFP^*; and (F/F’) *UAS-Axn^RNAi^/+; r4-Gal4/Abd-B^EGFP^*. Adipocytes with lower levels of Abd-A expression are marked with ‘>’ in E/E’, and adipocytes with low Wnt activity are marked with ‘*’ (asterisk) in F/F’ and H/H’. Scale bar in panel (H’): 10 µm.

**Supplementary Table S1. List of the *Drosophila* stocks used in this study.**

**Supplementary Table S2. Primers used in this study.**

## Data availability statement

The RNA-seq data in this study will be deposited in NCBI’s Gene Expression Omnibus upon acceptance of this work.

