## Supplementary figures and images for "Metabolic control by the Bithorax Complex-Wnt signaling crosstalk in *Drosophila*"

### Supplementary Figures S1-S10

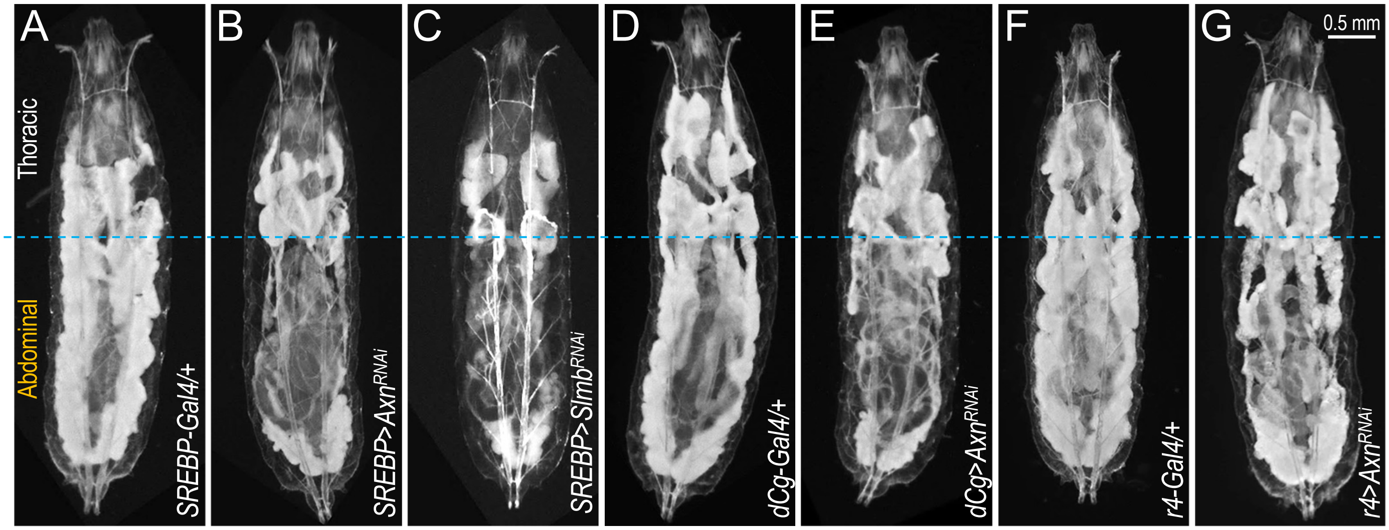


**Suppl Fig. S1**

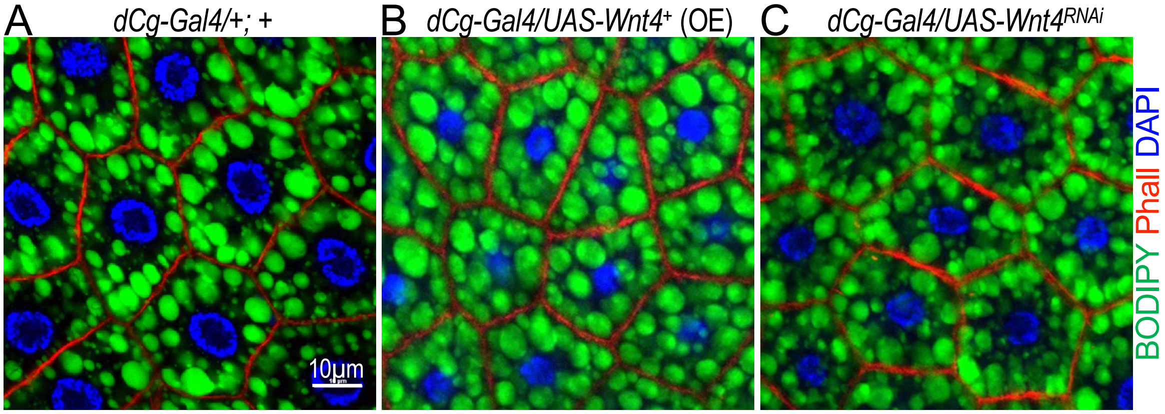


**Suppl Fig. S2**

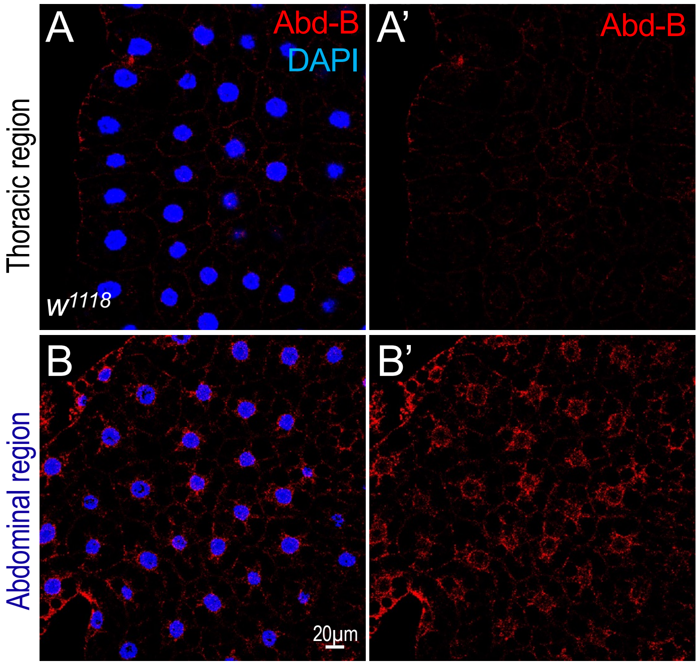


**Suppl Fig. S3**

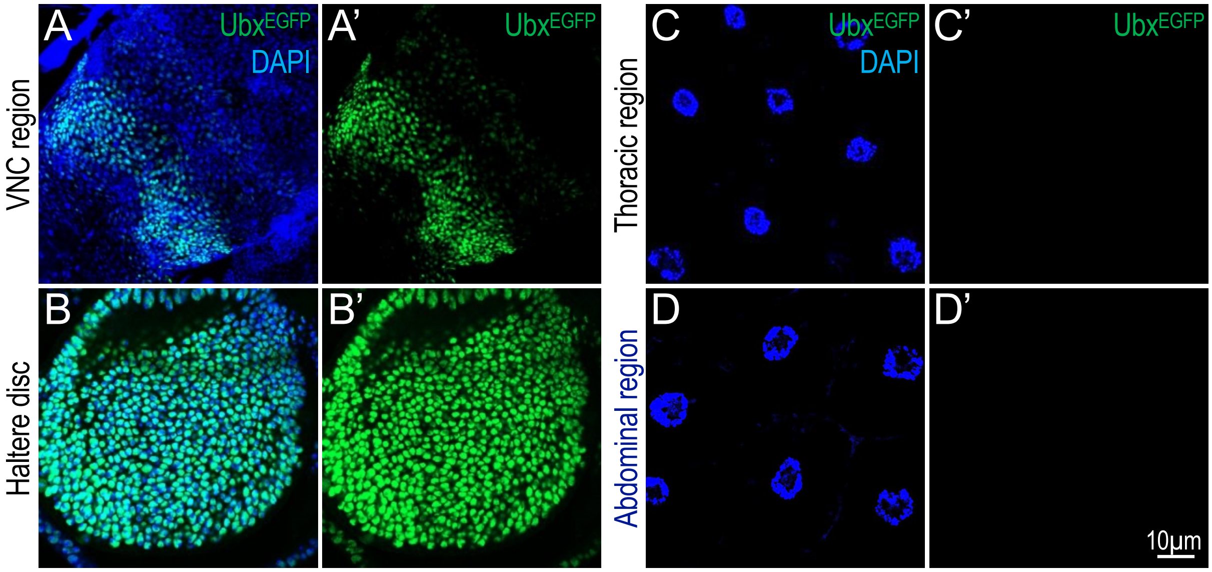


**Suppl Fig. S4**

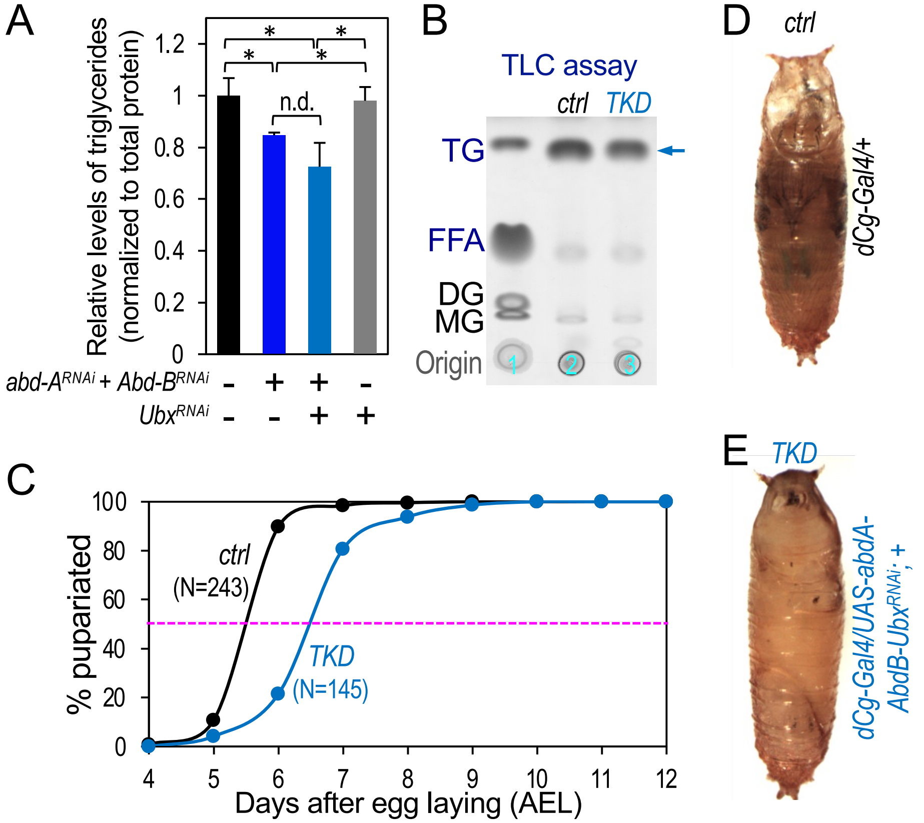


**Suppl Fig. S5**

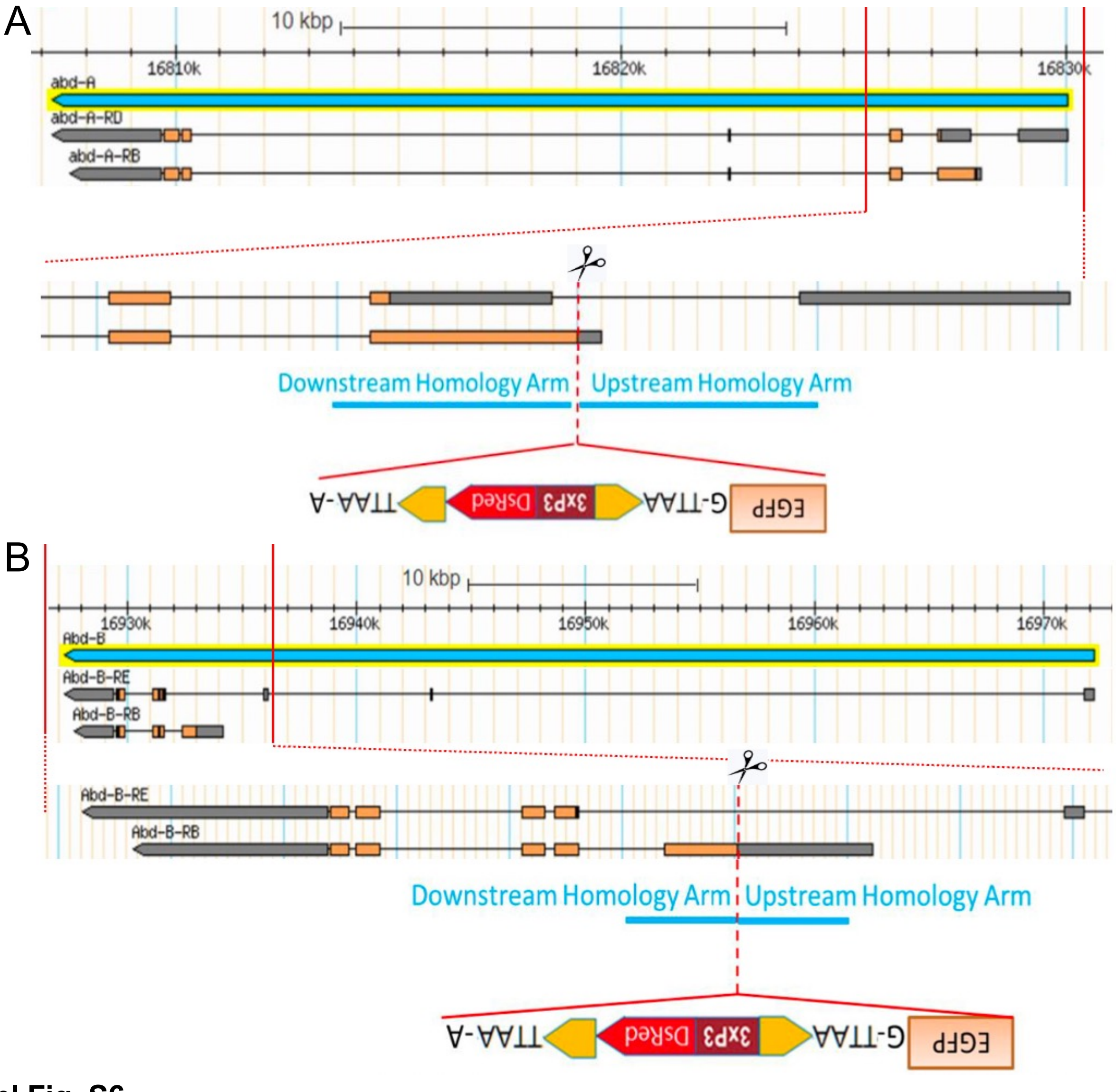


**Suppl Fig. S6**

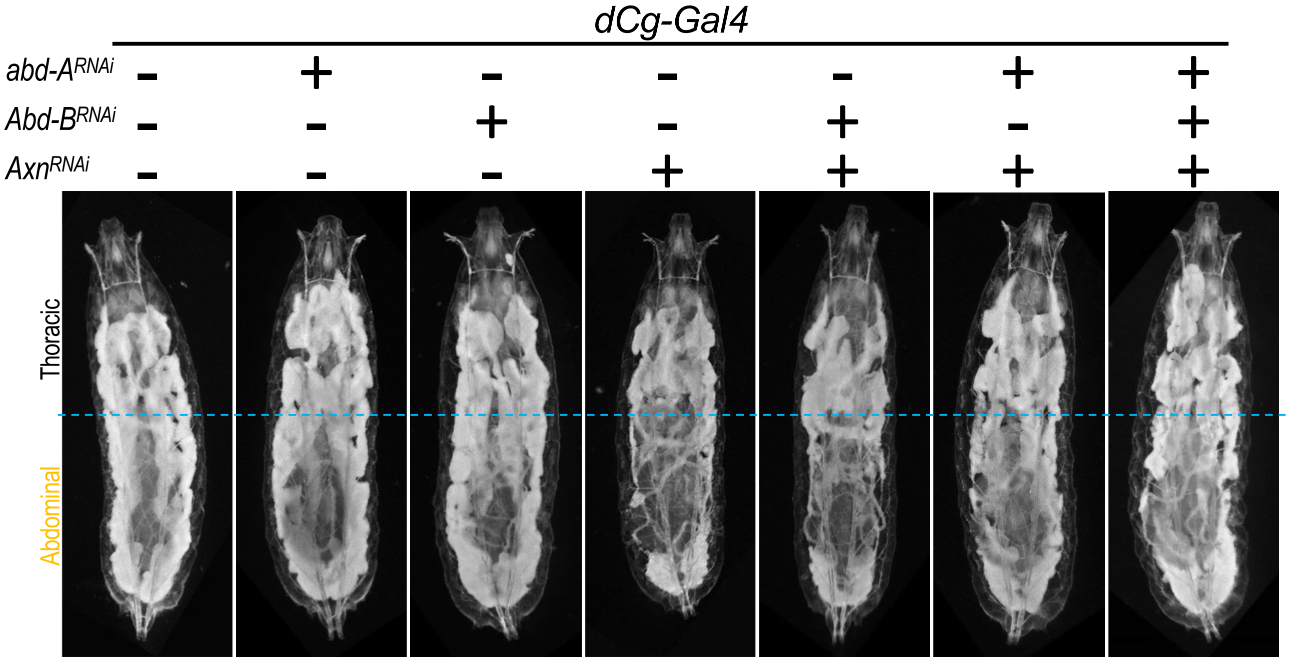


**Suppl Fig. S7**

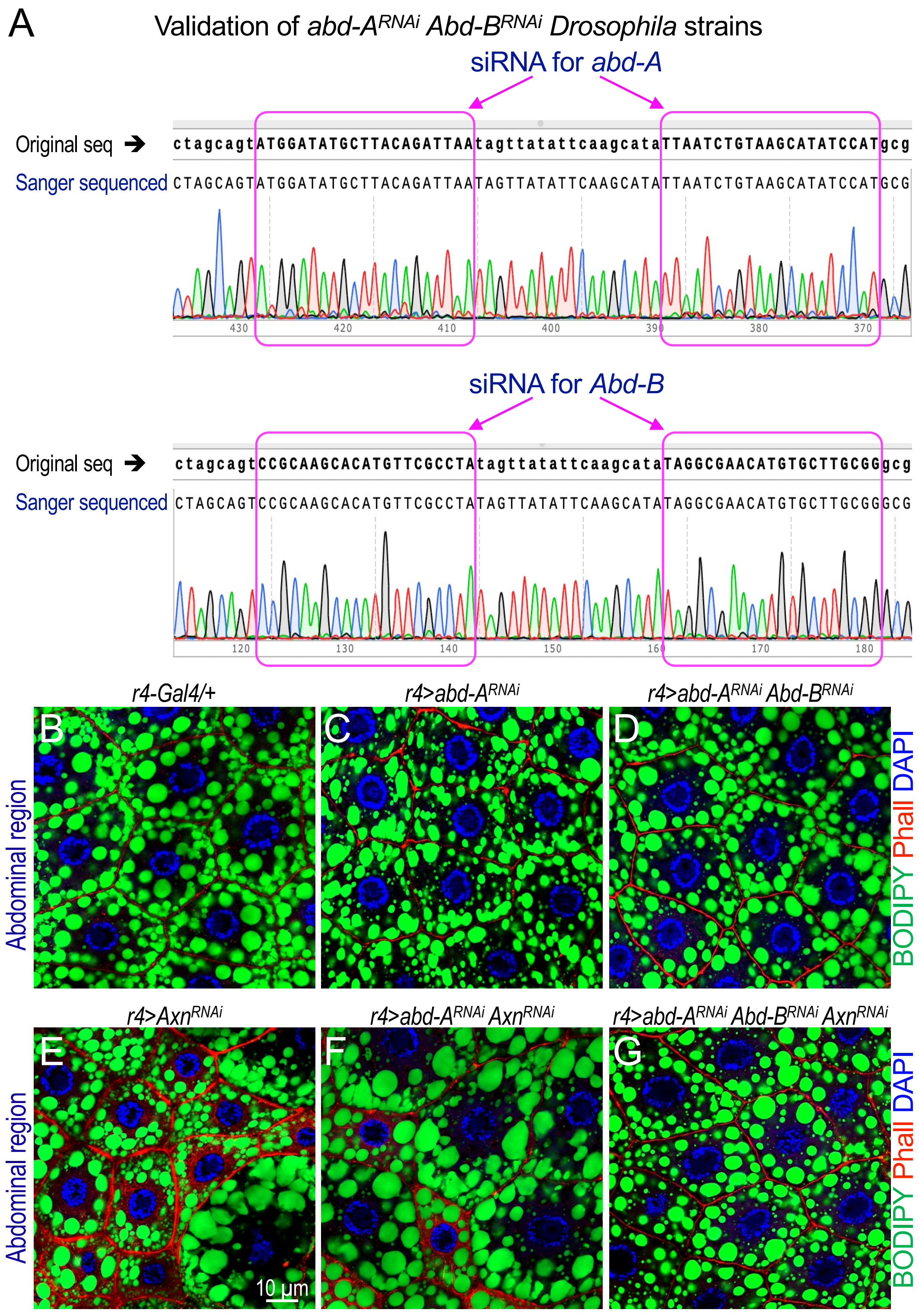

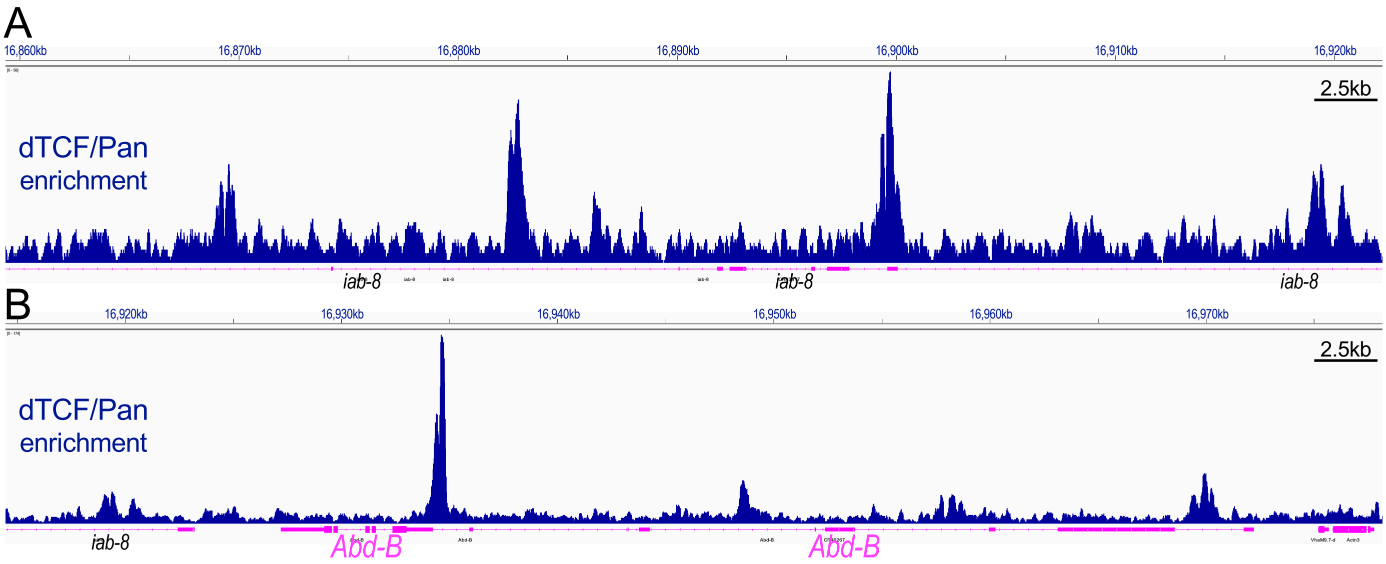

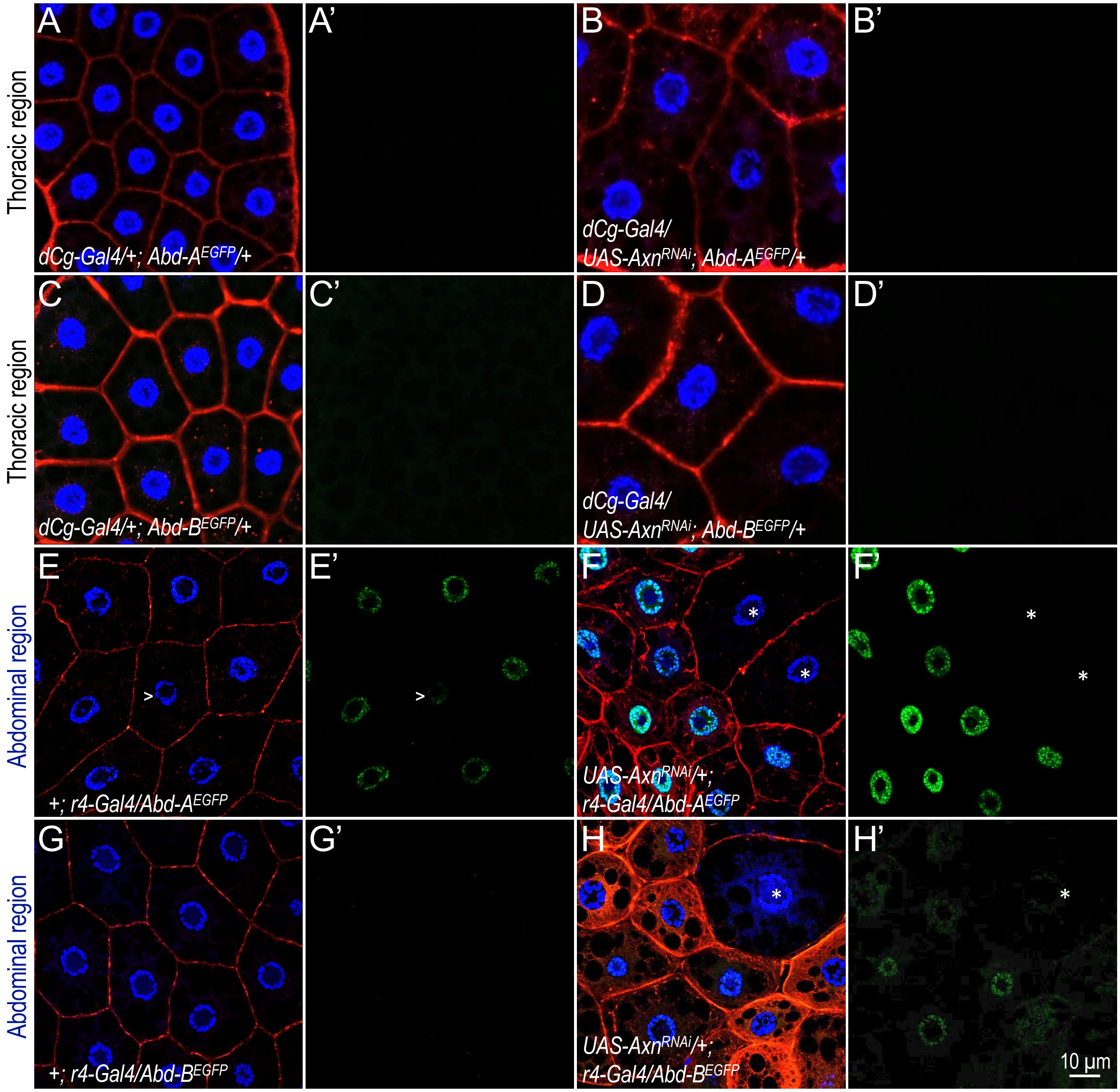


**Suppl Fig. S10**
